# Predictive feedback, early sensory representations and fast responses to predicted stimuli depend on NMDA receptors

**DOI:** 10.1101/2020.01.06.896589

**Authors:** Sounak Mohanta, Mohsen Afrasiabi, Cameron Casey, Sean Tanabe, Michelle J. Redinbaugh, Niranjan A. Kambi, Jessica M. Phillips, Daniel Polyakov, William Filbey, Joseph L. Austerweil, Robert D. Sanders, Yuri B. Saalmann

## Abstract

Learned associations between stimuli allow us to model the world and make predictions, crucial for efficient behavior; e.g., hearing a siren, we expect to see an ambulance and quickly make way. While there are theoretical and computational frameworks for prediction, the circuit and receptor-level mechanisms are unclear. Using high-density EEG, Bayesian modeling and machine learning, we show that inferred “causal” relationships between stimuli and frontal alpha activity account for reaction times (a proxy for predictions) on a trial-by-trial basis in an audio-visual delayed match-to-sample task which elicited predictions. Predictive beta feedback activated sensory representations in advance of predicted stimuli. Low-dose ketamine, a NMDA receptor blocker – but not the control drug dexmedetomidine – perturbed behavioral indices of predictions, their representation in higher-order cortex, feedback to posterior cortex and pre-activation of sensory templates in higher-order sensory cortex. This study suggests predictions depend on alpha activity in higher-order cortex, beta feedback and NMDA receptors, and ketamine blocks access to learned predictive information.

## Introduction

The classical view of sensory processing focuses on feedforward information transmission from the sensory organs to higher-order cortex, to generate representations of the world [1, 2]. However, growing evidence of expectations strongly influencing perception and behavior [3–5] suggests that the brain actively predicts incoming sensory information, a process that is not featured in the traditional framework. Predictive coding (PC) takes this process into account, wherein the brain uses generative models to make inferences about the world [6–10], possibly even to support conscious experience [11–13]. PC proposes that these models, based on prior sensory experiences, are represented at higher-order levels of a cortical hierarchy. The model predictions are transmitted from higher-order to lower-order cortex along feedback connections. Any mismatch between feedback predictions and observed sensory evidence generates an error signal, which is transmitted along feedforward connections, to update models in higher-order cortex [14–16]. The updated model is used in the next iteration to generate new predictions. This process of Bayesian updating aims to optimize beliefs about the sensory world. However, the neural representation of predictions is unclear.

N-methyl-D-aspartate receptors (NMDARs) may play a key role in PC. Theoretical work on PC [17] has proposed that higher levels of a cortical hierarchy transmit top-down predictions to lower levels through NMDAR-mediated signaling. Consistent with this proposal, NMDARs have been shown to modulate higher-order (frontal) cortical excitability [18–21], be enriched in superficial and deep cortical layers where feedback connections terminate [22], and contribute to feedback activity [23]. However, there is a lack of experimental evidence linking NMDARs and prediction itself. Ketamine, a NMDAR blocker [24], can reduce prediction error signals, measured as auditory mismatch negativity (MMN) in oddball paradigms [25–27]. But because the MMN reflects the mismatch between predictions and observed sensory evidence, it is difficult to dissect if ketamine’s effect on the MMN is due to ketamine influencing feedback predictions, feedforward sensory evidence or error signaling directly, any of which can reduce the MMN. Further, prediction is only assumed in oddball paradigms; there is no behavioral measure of prediction. Hence, a paradigm that incorporates a behavioral readout of predictions and separates predictions from other PC mechanisms is required to probe a contribution of NMDARs to predictions and their neural representation.

To test circuit and receptor-level mechanisms of prediction, we recorded 256-channel EEG of subjects performing an audio-visual delayed match-to-sample task (Fig 1A). The task design temporally separates predictions (generated during the delay period) from error processing (after image onset), which cannot be done in oddball paradigms [25–27]. Furthermore, having auditory stimuli carry predictive information about visual stimuli allows us to modulate separate feedforward (auditory to frontal) and feedback (frontal to visual) pathways. Subjects performed the task before, during, and after recovery from sub-hypnotic dosing of ketamine, targeted to concentrations that modulate NMDARs. In control experiments, we instead administered sub-hypnotic dexmedetomidine (DEX), an α_2_ adrenergic receptor agonist, selected to account for changes in arousal and modulation of hyperpolarization-activated cyclic nucleotide channels (HCN-1, which mediate ketamine’s anesthetic effects [28]). We found that subjects responded most quickly to highly predictive sounds, reaction times (RTs) thus serving as a behavioral readout of predictions. Drift diffusion modeling [29], which links decision-making processes to RTs, demonstrated that both inferred “causal” relationships between stimuli and frontal alpha power predicted RTs on a trial-by-trial basis – here causal used in a statistical sense, i.e., subjects tracked how often each image was preceded by its paired as well as unpaired sounds – and these relationships were notably absent under ketamine. Frontal alpha power contributed most to accuracy when classifying predictive stimuli using a support vector machine, consistent with the proposal that predictions increase the signal-to-noise ratio (SNR) in frontal cortex. Moreover, frontal cortex transmitted predictions to posterior cortex using beta frequencies, measured as Granger causal influence, activating sensory representations of predicted images before their presentation. Because ketamine, but not DEX, blocked these neural representations of predictions and their behavioral advantage, this study suggests a key role for frontal and posterior alpha activity, beta-mediated feedback and NMDARs in prediction mechanisms, as postulated in PC frameworks.

**Fig 1.**
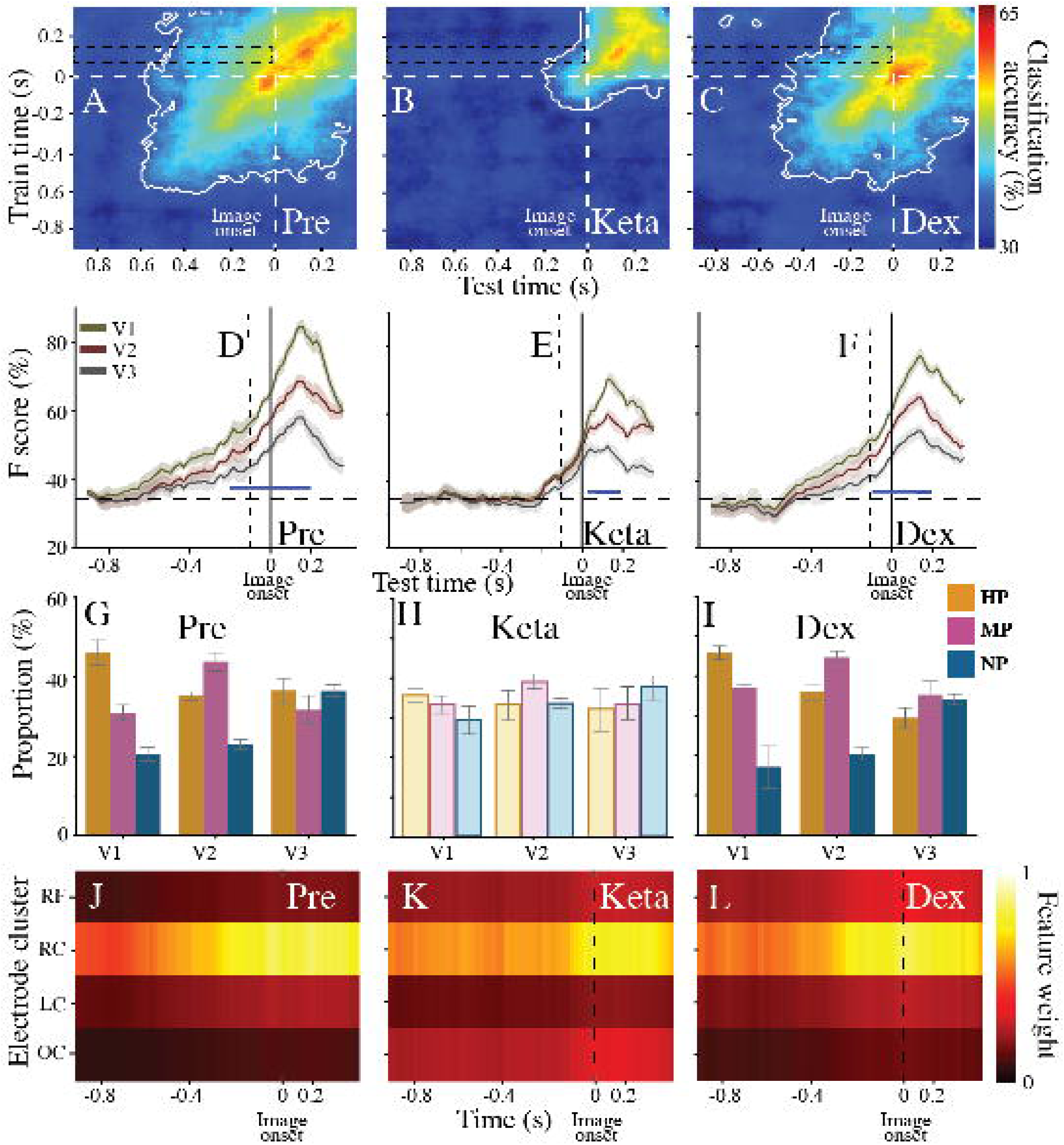
Ketamine blocked fast RTs to predictive sounds. See also Fig S1 and S2. (A) We manipulated subjects’ predictions by changing the probability of an image appearing after its associated sound in an audio-visual delayed match-to-sample task (after the initial learning phase). Population RT (+SE) of 15 subjects (B) before (Pre), (C) under (Keta), and (D) after recovery (Reco) from ketamine. (E) Population RT (+SE) of 14 subjects under dexmedetomidine (DEX). RTs from correct match trials. (F) Population causal power (CP) values across time for pre-ketamine (Pre) testing. Each data point corresponds to the CP at a particular trial for an individual subject. The data points form a trajectory of CP across trials. Data from 15 subjects overlaid. (G) HDDM of RTs. Bias (z) calculated for each trial (t) using CP. β_1_ determines the relationship between z and CP. (H) Posterior probability density of β_1_ for different drug conditions.

## Results

### Predictions improved RTs

Subjects initially learned paired associations (A1-V1, A2-V2, A3-V3) between three sounds (A1, A2, A3) and three images (V1, V2, V3) through trial-and-error. During learning, each sound and image had equal probability (33%) of appearing in any given trial, preventing subjects developing any differential predictions due to stimulus frequency. Thus, the presence of any given sound does not predict the occurrence of any future image, during this learning phase. Following the presentation of both stimuli, subjects reported if the sound and image were in fact paired, i.e., whether or not they matched. To manipulate subjects’ predictions during subsequent testing, we varied the probability of an image appearing after its associated sound. This probability was different for each sound: 85% chance of V1 after A1; 50% chance of V2 after A2; and 33% chance of V3 after A3 (Fig 1A). Thus, A1 was highly predictive (HP), A2 was moderately predictive (MP), and A3 was not match predictive (NP).

We hypothesized that increasing the predictive value of the sound would allow subjects to better predict the upcoming image, enabling quicker responses (HP<MP<NP) in match trials. However, if predictions are mediated by NMDARs, then ketamine should disrupt predictions; i.e., subjects administered with a sub-anesthetic dose of ketamine should be unable to exploit the differential predictive value of each sound, thus preventing faster RTs. If these effects are specific to NMDAR manipulation, then the control drug DEX should still allow faster RTs to predicted stimuli. To investigate these hypotheses (and to restrict multiple tests on the same dataset), we ran a linear mixed effects (LME) model. We used the sounds’ predictive value (HP, MP, NP) and drug condition (before drug, under ketamine, under DEX, after recovery from drug) as independent variables and RT as our dependent variable (RT∼ sounds’ predictive value + drug condition + sounds’ predictive value x drug condition). We applied a contrast analysis strategy to model our independent variables. Our study conformed to the guidelines set out by Ableson and Prentice [30] with regards to contrast analysis; i.e., we included the contrast of interest along with paired, orthogonal contrasts. (Contrasts of interest only explain a part of the total variation between groups. We included orthogonal contrasts to explain the residual variance. According to Abelson and Prentice [30], an analysis of the residual variance, i.e., orthogonal contrasts, is important since one may miss systematic patterns in the data if one only tests the contrast of interest. They suggested that finding a significant contrast of interest and a non-significant orthogonal contrast confirms the data support the hypothesis.)

To test our hypothesis that a greater predictive value of sounds allows subjects to respond faster, we used a linear contrast (HP, MP, NP: -1, 0, 1) as our contrast of interest, and a quadratic contrast (HP, MP, NP: -1, 2, -1) as the orthogonal contrast in the analysis. Further, to test our hypothesis that only ketamine prevents these faster responses, we used a four-level contrast (under ketamine, before drug, under DEX, after recovery: 3, -1, -1, -1) as our contrast of interest, and two orthogonal contrasts (under ketamine, before drug, under DEX, after recovery: 0, 0, -1, -1 and 0, 2, -1, -1) in the analysis. A significant main effect of prediction will confirm predictive sounds produce faster responses. A significant interaction effect of prediction and drug condition will confirm that ketamine disrupts subjects’ ability to exploit predictive sounds to respond faster.

We found a significant main effect of sounds’ predictive value in our LME model (ANOVA, F(1, 21.07)=14.14, P=0.001; orthogonal contrasts non-significant). RTs were faster when sounds had greater predictive value (Fig 1B). This result was further validated in parallel psychophysics experiments, where we controlled for possible match bias [31] using randomly interleaved “inversion trials” (in which subjects simply indicated whether greebles were inverted) to minimize the expectation of match trials (ANOVA, F(1, 21.92)=18.71, P=0.0001, orthogonal contrasts non-significant, Fig S1A). Furthermore, we found effects on RTs could not be explained by speed-accuracy trade-offs, as subjects were most accurate for HP, followed by MP and NP sounds (ANOVA, F(1, 21.66)=14.42, P=0.001, orthogonal contrasts non-significant, Fig S1B).

### Ketamine blocked fast RTs to predictive sounds

Importantly, we found a significant interaction of sounds’ predictive value and drug condition (ANOVA, F(1,16.42)=5.51, P=0.03). The interaction effect confirmed that, under ketamine, the linear correlation between the predictive value of sounds and RT was diminished (Fig 1C). This effect was specific to NMDAR manipulation, as DEX did not disrupt the ability of subjects to exploit the differential predictive value of the sounds. Under DEX, the linear correlation between the predictive value of sounds and RT was intact (Fig 1E), similar to the pre-drug baseline condition. These pharmacological effects were not due to low accuracy as subjects’ average accuracy was similar across all three conditions (77.8% under ketamine, 85.7% without ketamine, and 81.0% under DEX; ANOVA, F(1, 23.932)=1.36, P=0.22). Neither were effects due to the level of sedation as subjects were more alert under ketamine than dexmedetomidine (under ketamine, average modified observer’s assessment of alertness/sedation (OAA/S) score of 4.85 compared to 3.33 under DEX (5, awake – 1, unresponsive); unpaired t-test, P=0.003). Significant interaction effects in our LME model also confirmed that the linear correlation of RTs with predictive strength returns after recovery from ketamine (2-4 hours after ending ketamine administration, depending upon subject’s recovery; Fig 1D). Overall, our results demonstrate that subjects used predictive information to enhance behavioral performance and the NMDAR-blocker ketamine prevented this behavioral advantage.

### Subjects based predictions on inferred “causal” relationships between stimuli

We next investigated what information in the trial history subjects base predictions on; in other words, how do subjects learn the predictive relationship between stimuli. We tested two possibilities: (i) did subjects generate predictions by keeping track of simple co-occurrences of each sound-image pair, i.e., were they basing predictions on correlations; or (ii) did subjects not only track the occurrence of each image with its paired sound but also unpaired sounds, i.e., were they basing predictions on “causation”. Here, the term causation is used in a statistical sense, where subjects can infer a causal relationship between the initial sound and the image that closely follows in time. To clarify the difference between correlation versus causation in this context, let us consider a hypothetical example (adapted from [32]) in which you would like to test whether your new automatic sprinkler system worked overnight. In the morning, you walk outside and see that the grass is wet. Hence, you might think that the sprinkler operated overnight. In this case, your inference is based on the correlation of two events. However, after learning from the weather report that it rained last night, you lose confidence that your sprinkler watered the lawn, i.e., the wet grass may be due to the rain instead. Here, the presence of one cause (rain) casts doubt on the other (sprinkler) and thus helps us develop a more appropriate cause-effect relationship between events (unlike simple correlation). To relate this back to our task, when subjects track the occurrence of each image with its paired sound but also unpaired sounds, it is similar to checking the weather report to deduce if the sprinkler worked properly. Inferring a cause-effect structure between stimuli helps subjects eliminate weak/conditional relations between particular sounds and images. To measure correlation (option i above), we calculated the transitional probability, i.e., how often a particular image follows the sound only. To measure causation (option ii), we calculated the causal power [33] of the sound-image association, which is the amount of evidence that a sound “causes” a particular image, as opposed to a random different cause (Table S1 and methods show calculation details; we also calculated two other measures of “causation” – ΔP and causal support – and observed similar results). We updated the transitional probability and causal power estimates each trial, to account for the additional information available, i.e., the transitional probability/causal power value was the same for each stimulus at the start of the pre-drug baseline, but these values eventually systematically differed between stimuli as more trials were performed, reflecting the accumulating information from the trial history (Fig 1F, Fig S1E-F shows the causal power differentiating all stimuli earlier than transitional probability).

We next determined whether transitional probability and/or causal power can account for the behavioral results (RTs). To this end, we modeled subjects’ decision-making process using a drift-diffusion model (DDM). Evidence accumulates (drift process) from a starting point to one of two boundaries. Here, the boundaries represent the two possible outcomes for match trials only (correct and incorrect). The drift process stops when it reaches a boundary, indicating the choice, and the time taken to reach the boundary represents the RT for the trial (Fig 1G). The starting point of the drift process – here modeled from image onset – may be biased towards one of the boundaries, and it is determined by a bias parameter, z. This parameter represents the predictive value of the sound, which can be based on the transitional probability or causal power. A drift process that starts with a larger bias will reach the decision boundary quicker, resulting in a faster RT, i.e., more predictive sounds generate a larger bias and faster RT. Thus, whichever of transitional probability or causal power (through the bias parameter, z) yield better correspondence with subjects’ RTs will be the better indicator of the information subjects used to generate predictions.

To test this, we used hierarchical Bayesian parameter estimation (HDDM) [29], which calculates the posterior probability density of the diffusion parameters generating the RTs for the entire group of subjects simultaneously, while allowing for individual differences. We estimated the regression coefficients to determine the relationship between trial-to-trial transitional probability/causal power and biases estimated from the posterior predictive distribution. In other words, we calculated the bias for each trial that best predicted the RT. But, for each trial, the bias was constrained to depend on the transitional probability or causal power (equation, Fig 1G). Hence, for each trial we calculated the relationship (regression coefficient β_1_) between the bias and transitional probability/causal power that best predicted RT. Specifically, we estimated the posterior probability density of the regression coefficient (β_1_; Fig 1G) to determine the relationship between the bias and either transitional probability or causal power. We found causal power (deviance information criterion, DIC=-3197) predicted RTs better than transitional probabilities (DIC=-1795), i.e., causal power better captured the basis of prediction generation (option ii above). Further, bias was positively correlated with causal power (P{β_1_>0}=0.04; Fig 1H). This suggests that subjects based predictions on trial-by-trial updates of inferred “causal” relationships between sounds and images, rather than just correlations.

The HDDM also provides a framework to model drug effects. Thus, we repeated the above analysis of subjects’ behavior under ketamine and under DEX. If ketamine prevents predictive information from conferring a behavioral advantage, all sounds will generate similar biases; i.e., there will be no correlation between the bias and the predictive value of sounds, so β_1_ will be zero. Indeed, under ketamine, β_1_ was not different from zero (P{β_1_>0}=0.76; Fig 1H). In contrast, under DEX, bias positively correlated with the predictive values of sounds (all sounds generating equal vs different biases, P{β_1_>0}<0.00001; Fig 1H) similar to baseline. After recovery from ketamine, once again β_1_ was greater than zero (P{β_1_>0}<0.00001; Fig 1H) confirming that subjects had again generated larger bias for more predictive sounds. The question is: (a) did subjects re-gain access to previously learned and stored predictive information (Fig S2B) or (b) did they re-learn the predictive value of each sound after recovery from ketamine (Fig S2C)? To answer this, we used the HDDM to analyze the first 30 trials for each subject after recovery (translating to approximately 10 trials for each sound cue, for every subject). We found that, only for the former (option (a)), bias positively correlated with causal power (P{β_1_>0}=0.03; Fig S2D). This suggests that ketamine did not produce a loss of previously learned predictive information, but rather ketamine prevented access to the predictive information.

### Strength of predictive information correlated with frontal alpha power

We next investigated the circuit-level mechanism of prediction. To show task-responsive EEG electrodes, we averaged the time-frequency response across all trials and all electrodes (this selection procedure does not bias towards particular predictions). This revealed a task-related increase in baseline-corrected alpha power (8-14Hz; Fig S3B) before drug administration. Four clusters of electrodes, right frontal (RF), right central (RC), left central (LC) and occipital (OC), showed significant modulation of delay period alpha power compared to baseline, irrespective of sound (Fig 3A). Considering alpha power as an index of neural excitability (reduced alpha indicating reduced inhibition/increased excitability) [34, 35], one might expect stronger predictions to be associated with lower alpha power, reflecting greater activation of prediction-encoding neurons. This was the case for the RF cluster. There was greater reduction of alpha power across the delay period after more predictive sounds (Fig 2A-C). We hypothesized that this differential alpha modulation should characterize the pre-drug baseline and DEX conditions, but not ketamine as it prevented predictive information from conferring a behavioral advantage. To test this hypothesis, similar to the RT analysis (except that EEG was not recorded after recovery from drug), we ran a LME model. We regressed delay period alpha power at the RF electrode cluster on sounds’ predictive value (HP, MP, NP) and drug condition (before drug, under ketamine, under DEX) [RF alpha power ∼ sounds’ predictive value + drug condtion + sounds’ predictive value x drug condition]. To test the effect of sounds’ predictive value, we used a linear contrast (HP, MP, NP: -1, 0, 1) as our contrast of interest, and a quadratic contrast (HP, MP, NP: -1, 2, -1) as the orthogonal contrast in the analysis. To test our hypothesis that the RF alpha power modulation before drug will change under ketamine but not under DEX, we used a quadratic contrast (before drug, under ketamine, under DEX: -1, 2, -1) as our contrast of interest, and a linear contrast (before drug, under ketamine, under DEX: -1, 0, 1) as the orthogonal contrast in the analysis (see below). The significant main effect of sounds’ predictive value confirmed stronger predictions correlated with lower delay period alpha power at the RF electrode cluster (ANOVA, F(1, 26.24)=4.82, P=0.0003, orthogonal contrast non-significant; Fig 2A-D; and Fig S3D-F). Hence, frontal alpha reflected predictions.

**Fig 2.**
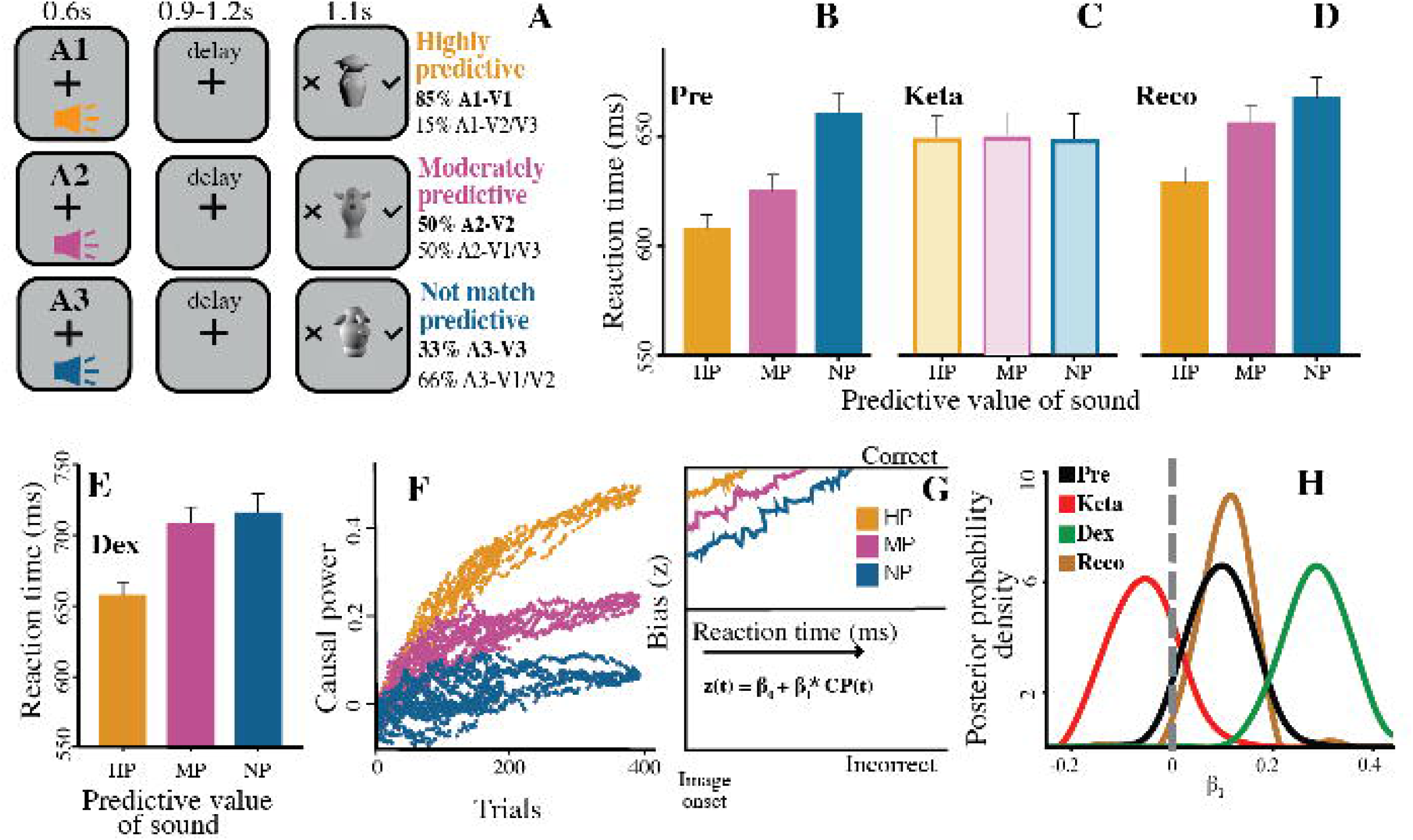
Right frontal alpha power correlated with prediction strength, and ketamine prevented this correlation. See also Fig S3. (A-C) Population time-frequency decomposition of right frontal electrode cluster (RF) before drug administration (Pre), for highly predictive (HP; A), moderately predictive (MP; B), and not-match predictive (NP; C) sounds. Power calculated in 0.55 s sliding windows, with window at 0 s representing interval -0.275 s to +0.275 s. (D-F) Population average RF alpha power in delay period (0.625 s to 0.275 s before image onset; this window size chosen based on wavelet length, to exclude stimulus-evoked activity – see methods) for (D) Pre, (E) ketamine (Keta) and (F) Dexmedetomidine (Dex).

**Fig 3.**
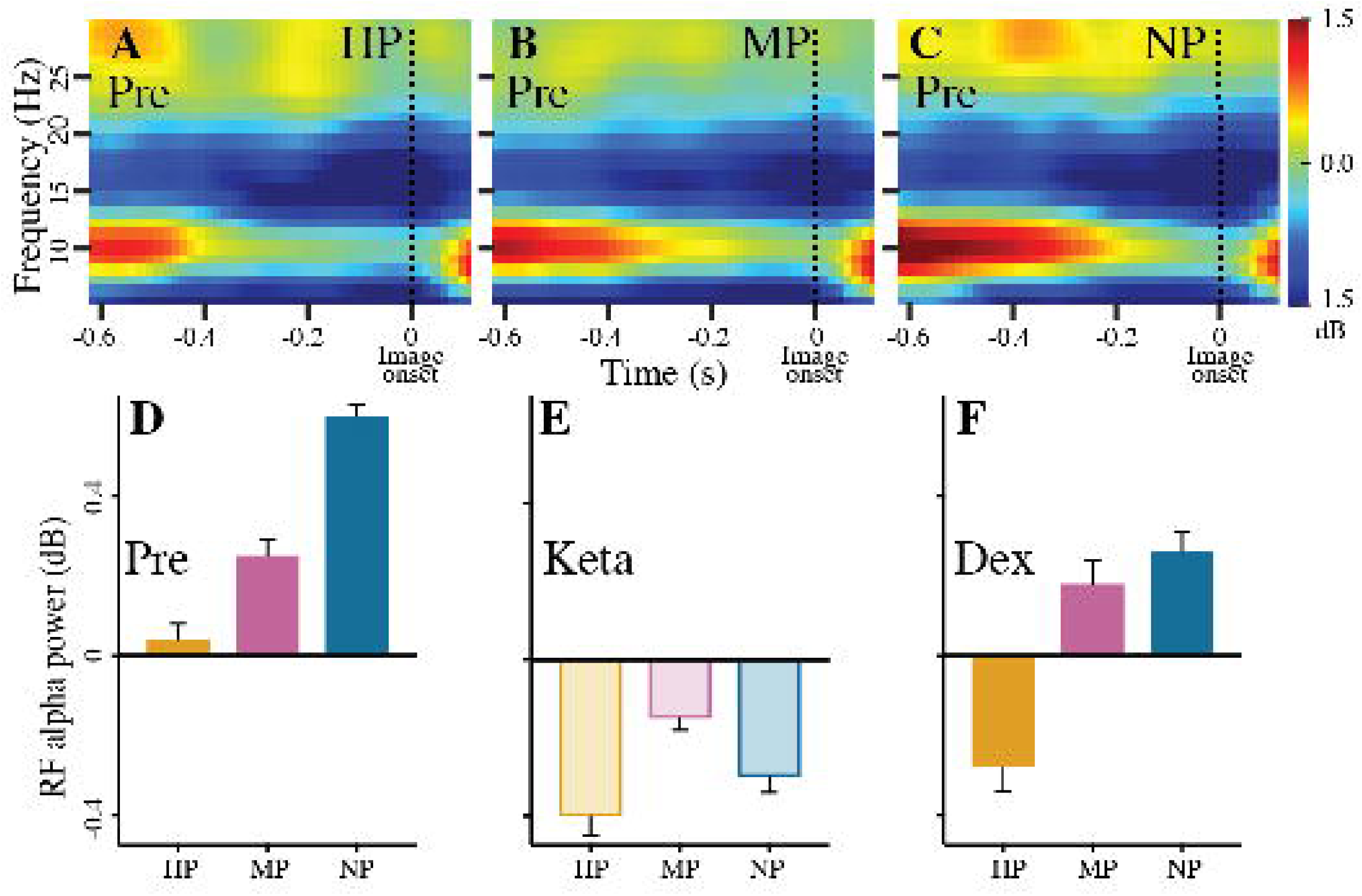
Ketamine prevented decoding of predictions. (A) Electrode clusters in right frontal (RF), right central (RC), left central (LC), and occipital (OC) cortex showing significant modulation in delay period alpha power (black circles represent the electrode that showed the greatest modulation, and its four nearest neighbors; we selected 5 electrodes in this way for consistency across clusters). Average across all trials (HP, MP and NP). (B-D) F-score using support vector machine (SVM) to decode predictive value (highly predictive, HP; moderately predictive, MP; or not-match predictive, NP) of sound, based on time-frequency power spectrum (B) before drug administration (Pre), (C) under ketamine (Keta), and (D) under dexmedetomidine (Dex). Dashed vertical line at 0 s denotes sound onset. Dashed vertical line at 1.225 s denotes earliest possible time window including image onset. Dashed horizontal line signifies level of decoding expected by chance. Gray shaded areas indicate significant zone of separation between HP, MP and NP, prior to image onset. (E) Posterior probability density of β_1α_. Bias (z) calculated for each trial (t) using alpha power (AP). β_1α_ determines relationship between z and AP. (F-H) Feature weights from SVM decoder for (F) Pre, (G) Keta, and (H) Dex. θ, α, β, and γ indicate theta (5-7 Hz), alpha (8-14 Hz), beta (15-30 Hz), and gamma (30-45 Hz) bands respectively.

To rule out the possibility that our RF spectral results are due to an impact on lower-level feedforward sensory processing, we also investigated if there were differences in the auditory ERPs of HP, MP and NP sounds. We tested for an effect over the entire time course between 50 to 300 ms after sound onset. Cluster-based permutation tests [36] revealed no significant difference between HP, MP and NP sounds in any electrodes at any latency. This confirmed that the correlation between the strength of predictions and frontal alpha power was not due to feedforward sensory processing, as all three sounds generated similar auditory ERPs (Fig S3A).

### Ketamine disrupted the correlation between predictions and frontal alpha power

NMDAR blockade has been shown to increase frontal cortical excitability [18–21], reducing response selectivity and SNR [37, 38]. We thus expect low-dose ketamine to increase frontal cortical excitability irrespective of the predictive value of sounds. This would manifest as similarly low RF alpha power for all sounds. Using our LME model, we found a significant interaction of prediction and drug condition (ANOVA, F(1,19.04)=2.81, P=0.0007). The interaction effect confirmed that, under ketamine, delay period alpha power at the RF electrode cluster was similar across sounds (Fig 2E). Whereas, for DEX, RF alpha power still showed a linear trend (Fig 2F) similar to the pre-drug baseline. This was not due to a more general drug-related change in alpha power, as power estimates were corrected based on the power prior to sound onset. Moreover, this was not due to a change in feedforward sensory processing under ketamine as cluster-based permutation tests [36] revealed no significant difference between HP, MP and NP sounds at any electrodes at any latency, i.e., all three sounds still generated similar auditory ERPs to that before ketamine. This suggests that ketamine affects mechanisms representing predictive information, and not basic sensorimotor mechanisms. That is, NMDARs mediate prediction strength through modulation of frontal excitability (reflected in alpha power).

### Decodability of predictions reduced under ketamine

Increased frontal excitability does not necessarily translate to useful prediction, if it reduces SNR (e.g., due to a more general change in excitability). We propose that increased frontal excitability (indicated by reduced alpha power) could facilitate better prediction, but this would not be the case if increased excitability reduced the SNR. To test this, we trained a decoder to measure if this hyperexcitability, reflected in low RF alpha power, still allows differential representation of predictions. Greater classification accuracy of each sound’s predictive value during the delay period would be consistent with a higher SNR of the neural representation of the prediction. Before ketamine, the classification F-score for each sound separated at 560 ms after sound onset (ANOVA, P=0.0018) and remained separate across the delay period until image presentation (Fig 3B). Similarly, classification F-score for DEX separated at 620 ms after sound onset (ANOVA, P=0.0067) and remained separate (Fig 3D). This shows that the more predictive the sound, the better the classification (HP>MP>NP). In contrast, under ketamine, there was no separation of classification F-score for each sound, and classification accuracy overall was lower (ANOVA, P=0.48; Fig 3C). This is consistent with ketamine disrupting the alpha indexing of predictive value. The weighting of features contributing to classifier performance confirmed that, before ketamine, RF alpha power contributed most to classification accuracy (RF alpha power feature (W_RFα_) > other features (W_∼RFα_), ANOVA, P=0.008; Fig 3F). This was also true under DEX (W_RFα_ > W_∼RFα_, ANOVA, P=0.013; Fig 3H). Under ketamine, RF alpha power contributed little to classification accuracy (W_RFα_ > W_∼RFα_, ANOVA, P=0.51; Fig 3G). These results suggest that ketamine disrupts the expression of predictive value in the power of RF alpha activity, which may be due to decreased SNR.

### Frontal alpha power correlated with RTs on trial-by-trial basis; but ketamine perturbed the correlation

Although frontal activity correlates with the predictive value of sounds, we need to show that subjects use it, i.e., that RF alpha power is linked to behavior. We again used the HDDM, but now to test whether RF alpha power predicts RT on a trial-by-trial basis. As before, the HDDM included two boundaries representing the possible choices in match trials, i.e., correct/incorrect. The starting point of the drift process – modeled from image onset – is determined by the RF alpha power in the delay period for each trial, which may be biased towards one of the boundaries, and is reflected in the bias parameter, *z*. We calculated the posterior probability density of the regression coefficient, β_1α_, which determines the relationship between the bias (z) and RF alpha power. Bias was inversely correlated with RF alpha power (P{β_1α_<0}=0.04, Fig 3E). This suggests that for more predictive sounds, lower RF alpha power creates a larger bias, and as a result, decisions are reached quicker (quicker RT). In contrast, ketamine blocked the correlation between bias and alpha power (regression coefficient, β_1α_, did not differ from zero, P{β_1α_<0}=0.51; Fig 3E). This suggests that subjects used frontal activity to make predictions.

### Predictive sounds activated sensory representations prior to visual stimuli; but ketamine perturbed these pre-stimulus activations

Prior work suggests that predictions can activate sensory templates prior to visual stimuli onset [39, 40]. To test which neural circuits and receptors contribute to this, we used a time generalized deep recurrent neural network (RNN) to classify visual stimuli (V1, V2, V3) at successive time bins (20 ms) across a trial. Visual representations are often activated as early as 50ms after stimulus onset [40]. To capture this early representation in our analysis, we trained our classifier with EEG time series data (not power) from the RF, RC, LC and OC electrode clusters, as spectral data corresponding to the visual stimulus presentation may be contaminated with pre-stimulus or motor activity due to the wavelet window size. Fig 4 A-C shows the cross-temporal decoding accuracy at different trained (y-axis) and tested (x-axis) time bins across the delay period and after visual stimulus onset. The white contour shows decoding accuracy significantly above chance (33%). We found significant classification accuracy above chance during the delay period well before visual stimulus onset for the “before drug” (Fig 4A) and DEX (Fig 4C) conditions. Crucially, when we trained the classifier on visual stimulus-evoked activity, e.g., at 100 ms post-stimulus onset (around the peak visual response) and tested in the delay period, as highlighted with the black dashed rectangle, there was significant decoding earlier in the delay period for the “before drug” and DEX conditions (pre-drug baseline: starting at 492 ms before stimulus onset; t(20,1000) =6.4, P=0.008; Fig 4A; and DEX: starting at 392 ms before stimulus onset; t(20,1000)=4.3, P=0.011; Fig 4C), compared to ketamine (starting at 116 ms before stimulus onset; t(20,1000) =2.23, P=0.03; Fig 4B). This confirmed activation of visual stimulus representations before stimulus onset; and that ketamine perturbed such pre-stimulus activations.

**Fig 4.**
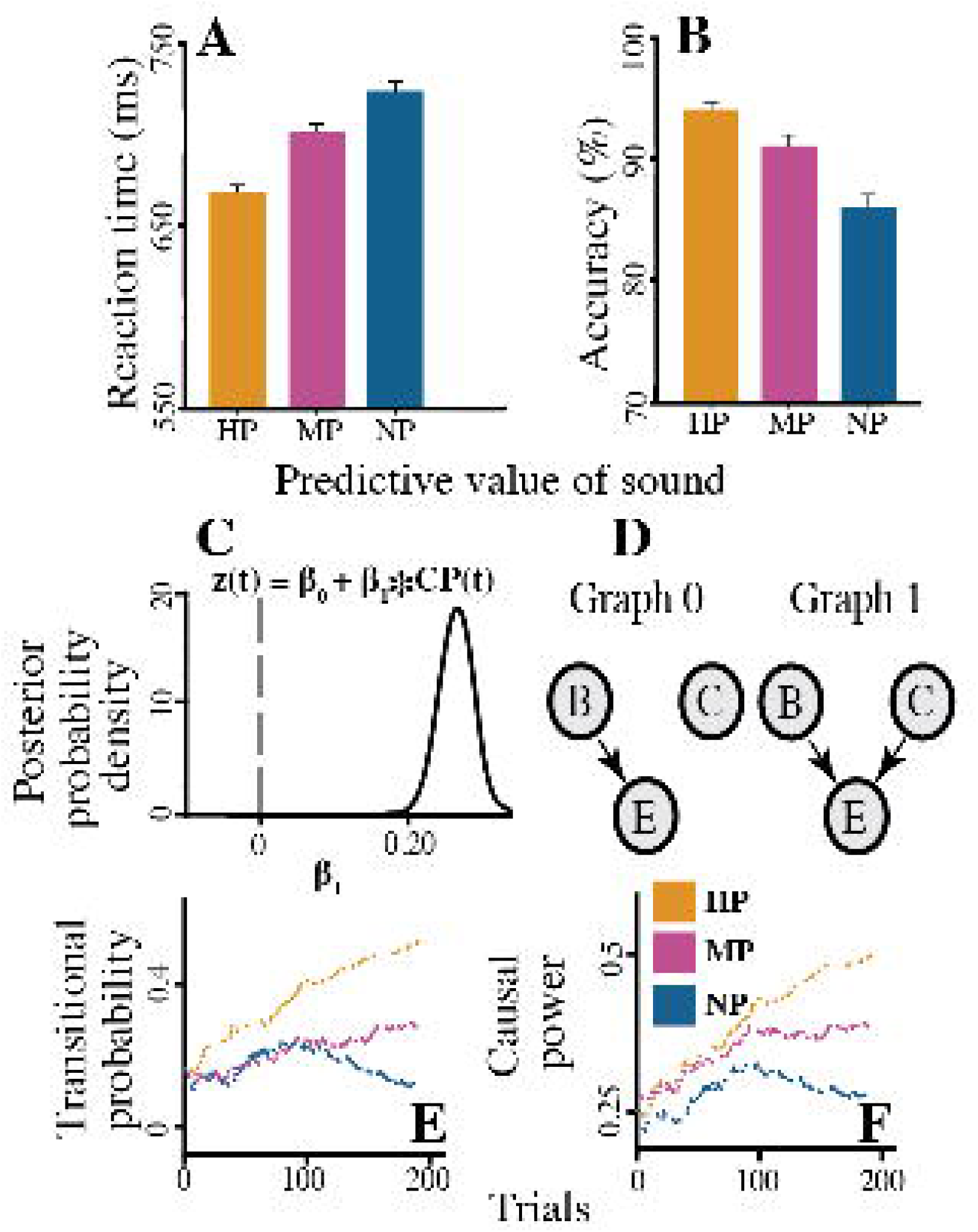
Ketamine perturbed pre-stimulus representations of predicted images. (A-C) Cross-temporal classification accuracy of images (V1, V2 or V3) based on time domain EEG data across all four clusters: (A) before drug administration (Pre), (B) under ketamine (Keta), and (C) under dexmedetomidine (Dex). Vertical and horizontal white dashed lines denote image onset. White solid contours in A-C show significantly above chance classification accuracy. Black dashed rectangle denotes time period used in D-F when classifier was trained on data 0.1 s post-visual stimulus onset and tested on the delay period pre-image onset. (D-F) Performance, measured as F-score, classifying V1, V2 and V3 trials when trained on 0.1 s post-image onset and tested on entire time period: (D) Pre, (E) Keta, and (F) Dex. Solid vertical black line denotes image onset. Dashed horizontal black line denotes chance level F-score. Dashed vertical black line denotes testing time, 0.1 s prior to visual stimulus onset, used in G-I. Blue horizontal line denotes significant difference between V1, V2 and V3’s classification. (G-I) Proportion of highly predictive (HP), moderately predictive (MP) and not-match predictive (NP) sound cues in trials correctly classified as V1, V2 and V3 by the classifier trained on 0.1 s post-visual stimulus onset and tested at 0.1 s before visual stimulus onset: (G) Pre, (H) Keta, and (I) Dex. (J-L) Feature weights when classifier trained at 0.1 s post-image onset and tested on the entire time across a trial in A-C: (J) Pre, (K) Keta, and (L) Dex.

Next, we probed which factors drive such pre-visual stimulus activations. We hypothesized that predictive sounds activate these early visual representations. That is, when subjects hear a predictive sound cue, they pre-activate a representation of its paired image during the delay period. Moreover, we posited that the strength of pre-activation will depend on the predictive value of the sound cue. Hence, the HP sound will pre-activate V1’s representation most strongly, followed by the MP sound moderately pre-activating V2’s representation, and the NP sound leading to little/no pre-activation of V3’s representation. To test our hypothesis, we measured how well the decoder classifies each of V1, V2 and V3 – expecting better classification performance for V1>V2>V3 – after training on visual-evoked activity (100 ms after stimulus onset reported here) and testing on the entire time across a trial. This relates to the row corresponding to the training time 100 ms post-visual stimulus onset in Fig 4A-C (black dashed rectangle), but now we further calculated the F-score for V1, V2 and V3 individually. During the delay period, we found the highest classification F-score for V1 followed by V2 then V3. For the “before drug” baseline, the classification F-score for V1, V2 and V3 separated at 200 ms prior to visual stimulus onset (ANOVA, F(20)=5.3, P=0.003) and remained separate thereafter, as shown by the blue horizontal line in Fig 4D. Similarly, the classification F-score for DEX separated 84 ms prior to visual stimulus onset (ANOVA, F(20)=3.2, P=0.011) and remained so (Fig 4F). In contrast, for the classification F-score under ketamine, there was no separation during the delay period, and the F-score curves only separated 12 ms after visual stimulus onset (ANOVA, F(20)=2.4, P=0.024; Fig 4E).

To further confirm that such differential pre-activation of visual representations are driven by the predictive value of their paired sound cues, we identified which sound cue (HP, MP or NP) preceded the correctly classified V1, V2 or V3 stimulus, when trained at 100 ms post-visual stimulus onset and tested at 100 ms prior to visual stimulus onset (vertical black dashed line in Fig 4D-F denotes testing time). One might expect HP sounds to precede pre-activation of their predicted V1 image representation, predominantly MP sounds to precede pre-activation of their predicted V2, and little bias for V3. For both “before drug” and DEX conditions, we indeed found that: for trials classified as V1, the proportion of HP sound cues was significantly higher than that for MP or NP (ANOVA, Pre-drug: F(20)=8.8, P=0.0001, Fig 4G; Dex: F(20)=4.3, P=0.001, Fig 4I); for trials classified as V2, the proportion of MP sound cues was significantly higher than that for HP or NP (ANOVA, Pre-drug: F(20)=6.1, P=0.0007, Fig 4G; DEX: F(20)=3.2, P=0.004, Fig 4I); but for trials classified as V3, there were no significant difference in the proportion of HP, MP and NP sound cues (ANOVA, Pre-drug: F(20)=0.80, P=0.07, Fig 4G; DEX: F(20)=0.66, P=0.1, Fig 4I). Conversely, under ketamine, there was no significant difference in the proportion of HP, MP and NP sound cues for any image (ANOVA, F(20)=0.33, P=0.31; Fig 4H).

Previous functional MRI [41] and EEG [42–44] studies found visual-evoked responses to greebles in higher-order sensory cortex of the right hemisphere. One might expect pre-activation of predicted greeble representations in the same region. Accordingly, the RC electrode cluster contributed most to the classification accuracy for both the “before drug” (ANOVA, F(20)=4.3, P=0.002; Fig 4J) and DEX (ANOVA, F(20)=3.8, P=0.01; Fig 4L) conditions, when we trained on data 100 ms post-visual stimulus onset and tested on the entire time across a trial. Crucially, the RC cluster significantly contributed to classification accuracy during the delay period for both “before drug” and DEX (476ms before visual stimulus onset for “before drug” and 344ms before stimulus onset for DEX). In contrast, under ketamine, the RC cluster did not significantly contribute to the classification accuracy until much later (52 ms before visual stimulus onset; ANOVA, F(20)=2.9, P=0.011; Fig 4K). Taken together, this suggests that NMDA receptors contribute to the pre-stimulus sensory templates activated by predictions.

### Strength of predictions correlated with feedback operating at beta frequencies

To investigate how predictions generated frontally influence pre-stimulus sensory templates, we measured functional connectivity between these frontal and sensory sites. Specifically, we calculated non-parametric spectral Granger causality in both EEG sensor and source spaces. For both spaces, we found frontal influences on sensory cortex dependent on the predictive value of sounds. We focus on source space results here (due to potential mixing of source signals in sensor space complicating connectivity analyses). Since frontal alpha power reflected the predictive value of sounds, we first calculated sources of sensor-level, task-related increases in baseline-corrected alpha power. ROIs in Fig 5A inset showed significantly increased source power during the delay period compared to baseline (p < 0.025, Bonferroni corrected across all cortical AAL ROIs, to ensure robust task-related sources). We found frontal cortical contributions to increased sensor-level alpha power in the right hemisphere only, as well as temporal cortical contributions consistent with previous greeble studies [41]. Consequently, we restricted our source-level Granger causality analysis between these sources within the right hemisphere. Using non-parametric spectral Granger causality, we measured the source-level feedback from the superior and medial frontal gyrus (Fig 5A, green) to inferior temporal gyrus (Fig 5A, red). Theoretically, the audio-visual task should anatomically isolate predictions transmitted along feedback pathways to posterior visual areas during the delay period from the feedforward auditory signals. One might expect higher frontal excitability (from stronger predictions or effect of ketamine) to give rise to stronger feedback. To test this, we ran the same LME model as for our previous EEG analyses. But here we regressed source-level beta band (15-30 Hz) feedback from the superior and medial frontal gyrus (Fig 5A, green) to inferior temporal gyrus on sounds’ predictive value (HP, MP, NP) and drug condition (before drug, under ketamine, under DEX) [beta feedback ∼ sounds’ predictive value + drug condition + sounds’ predictive value x drug condition]. To test the effect of sounds’ predictive value, we used a linear contrast (HP, MP, NP: -1, 0, 1) as our contrast of interest, and a quadratic contrast (HP, MP, NP: -1, 2, -1) as the orthogonal contrast in the analysis. To test our hypothesis that the beta feedback effect before drug will be perturbed under ketamine but not under DEX, we used a quadratic contrast (before drug, under ketamine, under DEX: -1, 2, -1) as our contrast of interest, and a linear contrast (before drug, under ketamine, under DEX: -1, 0, 1) as the orthogonal contrast in the analysis. A significant main effect of sounds’ predictive value (ANOVA, F(1,30.46)=11.47, P=0.0001, orthogonal contrast non-significant) showed stronger predictions were associated with greater Granger causal influence of right frontal cortex on right inferior temporal cortex in the beta band during the delay period (Fig 5, A and B). Taken together, this suggests predictions are disseminated along feedback connections down the cortical hierarchy prior to image onset.

**Fig 5.**
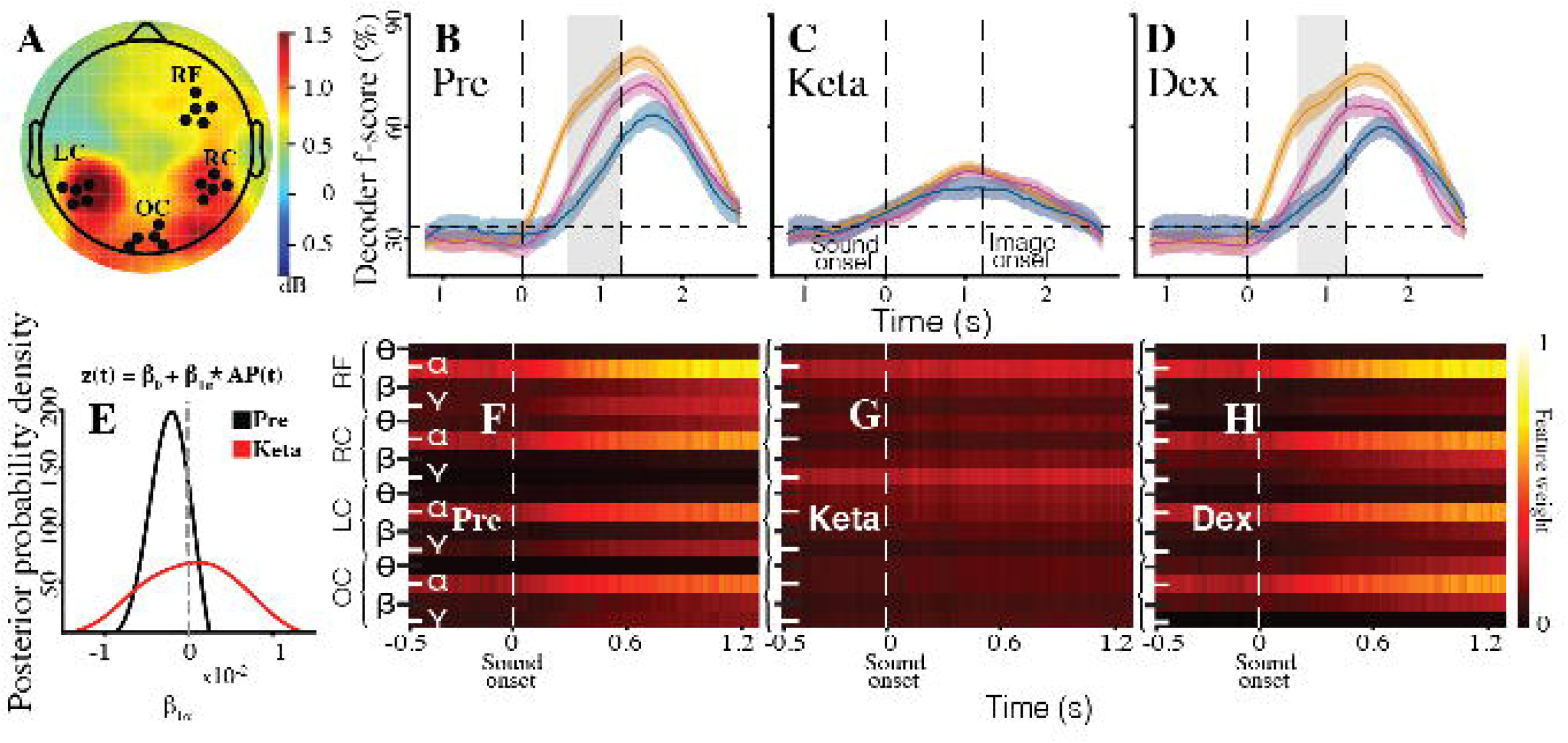
Beta feedback correlated with prediction strength, but ketamine perturbed this correlation. (A) Population average non-parametric Granger causal influence from right frontal to right inferior temporal cortex in the beta band (15-30 Hz) during the delay period (starting 0.9 s before image onset). Highly predictive, HP; moderately predictive, MP; not-match predictive, NP. Inset shows regions of significantly increased source power during delay period compared to baseline (P < 0.025, Bonferroni corrected across all cortical AAL regions) after beamforming source localization. Green, superior and middle frontal gyrus; Red, inferior temporal gyrus. (B) Population average non-parametric spectral Granger causal influence during delay period. (C) and (D) Population average non-parametric Granger causal influence from right frontal to right inferior temporal cortex in beta band during delay period for (C) ketamine (Keta) and (D) dexmedetomidine (Dex).

### Ketamine perturbed the correlation between prediction strength and beta feedback

Because NMDAR blockers have been reported to perturb feedback pathways in macaques [23] and humans [45], we expected ketamine to alter feedback (carrying predictions) from frontal to inferior temporal cortex. We do not expect such modulations of feedback pathways under DEX as DEX did not change correlation of RT and frontal alpha power with prediction. We found a significant interaction of prediction and drug condition (ANOVA, F(1,24.98)=2.61, P=0.04). The significant interaction effect demonstrated that, under ketamine, there was no longer a correlation between the predictive value of sounds and beta band Granger causal influence of right frontal on right inferior temporal cortex (Fig 5C); i.e., ketamine scrambled the feedback for each sound. In contrast, for DEX, the Granger causal influence in the beta band still differed between sounds (Fig 5D). This suggests that the predictive feedback facilitating behavior depends on NMDARs.

## Discussion

Our results show NMDAR-mediated, circuit-level mechanisms of prediction and its behavioral effects. Frontal cortex represented predictions and, starting prior to image onset, transmitted them to posterior cortex in the beta band, to activate a sensory representation of the predicted image. Stronger predictions enabled faster responses, and reflected causal power, i.e., inferred “causal” relationships between sounds and images. In contrast, ketamine prevented fast responses to predictive stimuli, as well as subjects from using the strength of causal power to generate predictions. At the circuit level, ketamine disrupted predictions by reducing frontal alpha power to the same low level prior to all images (likely indicating reduced SNR), leading to undifferentiated feedback and perturbed pre-stimulus sensory activations. Overall, it suggests that NMDARs normally sharpen representations of predictions in frontal and posterior cortex, to enable PC. The data are less supportive of the classical view of perception, with its emphasis on feedforward processing to reconstruct images because one might have expected little systematic difference in behavioral and neural measures for different predictive conditions.

The initial predictive auditory stimulus will activate auditory pathways, leading to the generation of a prediction of the subsequent visual image by a higher-order, multi-modal area. Complex auditory stimuli, like the trisyllabic greeble names in our task, are represented in auditory lateral belt and parabelt cortex [46]. Belt and parabelt regions are connected with a number of multi-modal areas in superior temporal and prefrontal cortex [47, 48], where there are multisensory neurons responding to both vocalizations and images [49, 50]. Recent work [51] suggests that early sensory fusion occurs in temporal or parietal multimodal areas, whereas more flexible weighting and integration of sensory signals for adaptive behavior occurs in frontal multimodal areas. Our results are consistent with this, but go further by showing that the frontal multimodal areas are the source of predictive multimodal signals, which, in addition to the enhancement of sensorimotor processing shown here, may be useful for communication and language processing [52]. Interestingly, we found that the frontal source and posterior target – showing pre-stimulus sensory activations of the predicted greeble – of predictive information was lateralized to the right hemisphere. This is consistent with previous studies suggesting a possible right hemispheric bias for the processing of greebles [41,53,54], and possibly faces more generally [55, 56].

There have been two broad approaches for understanding how we learn relationships between stimuli: associative and causal approaches. Classical theories like the Rescorla-Wagner model [57–59] propose learning as the association between a cue and an outcome. Causal models of learning [33,60,61], on the other hand, propose that we learn relationships between latent unobservable “causes” and observable stimuli (both cues and outcomes). In other words, causal learning models have put forward the idea of “clustering”, where observations (related to both cue and outcome) are clustered together according to their hypothetical latent causes. In line with previous work [62, 63], we found that subjects’ RTs were best predicted by a causal model (casual power) and not an associative model (transitional probability). Our causal learning results have further implications on models of sequential learning. Previous work on causal learning focused on summarized data contingency [33, 64]. Our findings support trial-by-trial learning from sequential data as proposed by a recent modelling study [65]. Additionally, using HDDM we found that both RF alpha power and causal power correlated to RT on a trial-by-trial basis. This points towards a neural readout of causal inference in humans.

Although beta oscillations have been proposed to maintain the current brain state [66, 67], there is growing evidence of beta activity playing a more dynamic role [68]. It has been proposed that beta oscillations are suitable for endogenous re-activation of cortical representations, to facilitate task-relevant activity patterns and cognitive demands [68]. In line with this, we found beta band predictive feedback from right frontal to inferior temporal cortices reactivates greeble representations in posterior cortex prior to image onset. Spitzer and Haegens [68] speculate that beta band activity is well suited to be a ‘transit’ between alpha frequency (generally associated with cortical excitation/inhibition [69, 70]) and gamma frequency (generally associated with population spiking and active stimulus coding [71]) activity. Our finding of frontal cortical alpha power coding for the predictive value of sound cues, followed by frontal influence on temporal areas (shown to have greeble representations [41,42,44]) at beta frequencies prior to visual-evoked activity, supports beta’s role as a ‘transit’ band. Intracortical laminar recordings in animal studies of sensory and attentional processing suggest that feedforward signaling operates at gamma frequencies, whereas feedback signaling operates at lower frequencies [72–74]. Consistent with this, previous work on predictive processing using univariate measures has reported the involvement of various lower frequencies, including theta, alpha and beta bands [75–77]. Although there are varying reports of the direction of alpha power changes in predictive processing, this may be due to “when” or “what” is being predicted, differences in task structure (e.g., analyzing the stimulus or delay period) and differences between brain regions [75, 78]. Further considering connectivity, a recent macaque study reported greater alpha and beta feedback from prefrontal to visual cortex during the presentation of more predictive visual stimuli [76]. Van Pelt et al. [79] also found strongest top-down feedback connectivity in the beta band while subjects viewed videos of predictable events. Our finding of frontal cortex causally influencing posterior cortex in the beta band according to predictions extends this finding to the delay period in the absence of sensory stimulation, as well as provides support for PC models that incorporate a key role for oscillatory activity more generally [75,80,81].

NMDARs are located in both superficial and deep cortical layers [22], potentially allowing NMDARs to modulate the representation of predictions in deep layers, as proposed in certain PC models [80], and predictive feedback signaling to superficial and/or deep layers. NMDAR blockade in humans has been shown to modulate frontal cortical excitability [20, 21], and our decoding analyses suggest that this NMDAR-related change in excitability reduces the SNR of prediction representations. This is consistent with macaque experiments showing that NMDAR blockade reduces the SNR in frontal cortex during the working memory period of an anti-saccade task [37]. In our study, NMDAR blockade also perturbed predictive feedback, consistent with macaque experiments showing NMDARs contribute to feedback signaling [23]. Taken together, these results suggest NMDARs influence both the representation of predictions in higher-order areas and predictive signaling to lower-order areas, which impacts the formation of pre-stimulus templates.

Ketamine at sub-hypnotic doses perturbed feedback connectivity from frontal to more posterior cortex – but not evoked activity in sensory cortices – during which subjects could still perceive and accurately respond to audio-visual stimuli. This raises questions about the requirement of frontal feedback integrity for consciousness. Further, it has been proposed that generative models create virtual realities that support conscious experience [11–13]. That subjects’ predictions in our study could be disrupted without impairing consciousness imposes constraints on PC as a theory of consciousness.

## Materials and Methods

### RESOURCE AVAILABILITY

#### Lead Contact and Materials Availability

This study did not generate new unique reagents. Further information and requests for resources, equipment and experimental methods should be directed to and will be fulfilled by the Lead Contact, Yuri Saalmann.

#### Data Availability Statement

Data generated in the study can be found here https://osf.io/3z49m/

### EXPERIMENTAL MODEL AND SUBJECT DETAILS

The University of Wisconsin-Madison Health Sciences and Social Sciences Institutional Review Boards (IRBs) approved experiments. 29 participants (14 female) performed the psychophysics predictive coding experiment. We excluded data from four subjects as their performance accuracy was below 50%. 17 additional participants (six female) for dexmedetomidine and 15 participants (five female) for ketamine took part in the pharmacology predictive coding experiments (mean age = 22.35 years; SD = 2.82 years). Of the 15 participants who performed ketamine experiments, the first 12 had to participate in the dexmedetomidine experiments first as per IRB requirements, whereas the final three participated in ketamine experiments first. The different orders of drug experiments yielded similar results. Three participants also performed in saline control experiments, which yielded results similar to the pre-drug baseline. Three participants (from 17 dexmedetomidine participants) were excluded for low accuracy (accuracy less than 50%). Participants who performed the psychophysics predictive coding experiment did not take part in the pharmacology experiments. We obtained informed consent from all participants.

### METHOD DETAILS

#### Stimuli

For our psychophysics experiments, we used biomorphic visual stimuli from Michael Tarr’s lab (http://wiki.cnbc.cmu.edu/Novel_Objects). These are known as greebles. Fig 1A shows examples presented to participants. We used three gray-scale greebles for each psychophysics session, and each greeble was personified with a name. We used novel sounds (trisyllabic nonsense words) for the greeble names – e.g., “Tilado”, “Paluti”, and “Kagotu” – from Saffran et al. [82]. The sounds were generated using the Damayanti voice in the “text to speech” platform of an Apple MacBook. To avoid differences in the salience of stimuli, greeble images have similar size (13 degrees of visual angle in height and 8 degrees in width), number of extensions and mean contrast, and greeble names have the same number of syllables and sound level (80 dB SPL).

For the pharmacology experiments, we generated three new triplets of greebles. To control for saliency, each participant rated greeble salience for each of four (three plus one from the psychophysics experiment) triplets, i.e., the participant identified whether any of the greebles in a triplet stand out compared to the other two greebles from the same triplet. We proceeded to use triplets which the participant rated all three greebles as being equally salient. We then named each of these greebles with a new trisyllabic nonsense word.

#### Audio-visual delayed match-to-sample task

Each trial of the task involves the sequential presentation of a sound (trisyllabic nonsense word) followed by a greeble image. We refer to stimuli using the following notation: A1, A2 and A3 correspond to each of the three sounds used (A for auditory) and V1, V2 and V3 correspond to each of the greebles used (V for visual). Using this notation, audio-visual stimulus sequences containing the matching name and greeble are A1-V1, A2-V2 and A3-V3. Audio-visual stimulus sequences containing a non-matching name and greeble are A1-V2, A1-V3, A2-V1, A2-V3, A3-V1 and A3-V2. We pseudo-randomized names for greebles (i.e., matching sounds and images) across subjects.

*Learning Phase.* During the first phase of the task, participants learn the association between the sounds and images (i.e., names of the greebles) through trial-and-error, by performing a match/non-match (M/NM) task. This phase is called the “learning” phase. Each trial starts with blank blue screen (R=35, G=117, B=208, 200ms duration; shown as gray in Fig 1A). After that, a black fixation cross (size 1.16 degrees of visual angle; jittered 200-400 ms) is presented followed by a sound, a greeble name voiced by the computer (600 ms duration). After a jittered delay period (900-1,200 ms duration), a greeble image (until a participant responds or 1,100 ms duration, whichever is earliest) was presented on the monitor screen, as well as two symbols (**√** and **X**) to the left and right of the greeble (9.3 degrees of visual angle from screen center). These symbols indicated participants’ two response options: match (√) or non-match (**X**). The symbol location, left or right of the greeble image, corresponded to the left or right response button, respectively: left and right arrow keys of a computer keyboard in the psychophysics experiments; and left and right buttons of a mouse in the pharmacology experiments. We randomly varied the symbols’ locations relative to the greeble image to minimize motor preparation (i.e., on some trials, a match response required a left button press and, on other trials, a match response required a right button press). In the learning phase, each greeble name and image had 33% probability of appearing in any given trial. This is to prevent subjects from developing any differential predictions about the greebles due to greeble name or image frequency, during the learning phase.

To address possible same-different biases, e.g., quicker reaction times (RTs) for match trials [31] we introduced a control called “inversion trials” in psychophysics experiments, to minimize the expectation of M/NM trials which, in itself, might otherwise contribute to participants’ responses. In these inversion trials, participants had to respond whether the greeble image presented on screen is inverted (the appropriate response button, left/right, indicated on screen by the left/right location of a red arrow pointing down) or upright (yellow arrow pointing up). Participants did not know the type of trial in advance; the trial type was only revealed by the symbols to the left and right of screen center at the onset of the greeble image (i.e., **√** and **X** signal M/NM trials, whereas downward red arrow and upward yellow arrow signal inversion trials). 50% of the total number of trials in the learning phase were inversion trials and the rest were M/NM trials. Because participants cannot specifically prepare in advance for M/NM trials due to the random presentation of inversion and M/NM trials, there should be minimal confounding of RTs with a bias towards match responses.

*Testing phase.* Once participants show above 80% accuracy for the M/NM trials in the learning phase of the task, they move on to the “testing phase” (1,000 trials for the psychophysics experiments; Fig 1A). During the testing phase, we manipulated predictions by changing the probability of a greeble appearing after its learnt name. This probability is different for each greeble name and image. That is, in the testing phase, when a participant hears A1, there is 85% chance of V1 being shown (highly predictive; HP); when a participant hears A2, there is 50% chance of V2 being shown (moderately predictive; MP); and when a participant hears A3, there is a 33% chance of V3 being shown (not-match predictive; NP). This allows participants to make stronger predictions about the identity of the upcoming visual image after hearing A1, than after hearing A2 or A3, for instance.

Inversion trials also consisted of half the total trials in the testing phase of the psychophysics experiments. We randomly presented all the trial types (M/NM trials and inversion trials) to the participants. The testing phase of the task had approximately equal match and non-match trials to avoid response bias. The testing phase also had an approximately equal number of trials for each greeble image, and its corresponding name had approximately equal probability of being voiced, to control for stimulus familiarity.

#### Causal Strength and Transitional Probability

We quantified the relationship between a sound (name) and its paired image (greeble) using the strength of causal induction: given a candidate cause *C* (sound) how likely is the effect *E* (i.e., how likely is it followed by its paired image). We will represent variables *C* and *E* with upper case letters, and their instantiations with lower case letters. Hence, *C* = *c* + / *E* = *e* + indicates that the cause/effect is present, and *C* = *c* – / *E* = *e* – indicates that the cause/effect is absent (for brevity, we will shorten variables equal to outcomes, such as *C = c +* or *C* = *c* – as simply *c* + or *c* –, respectively). The evidence for a relationship can be encoded as a 2 X 2 contingency table for each sound, as in Table S1 (black letters), where N(c+, e+) represents the number of trials in which the effect occurs in the presence of the cause, N(c–, e+) represents the number of trials in which the effect occurs in the absence of the cause and so on. Applied to our study, e.g., *C* could be hearing sound A1, and *E* viewing the paired greeble V1. For this case, N(c+, e+) would be the number of trials V1 follows A1; whereas N(c–, e+) would be the number of trials V1 follows A2 or A3. The full contingency table for the “Highly Predictive” auditory cue A1 and its paired greeble V1 is shown in Table S1 (in green letters). There are analogous contingency tables for the other two auditory cues and their paired greebles.

Based on these contingency tables, we calculated three different measures of causal relationship (ΔP, causal power and causal support) for each trial. ΔP and causal power assume that *C* causes *E*. ΔP reflects how the probability of *E* changes as a consequence of the occurrence of the cause *C*. Causal power corresponds to the probability that an effect *E* happened because of cause *C* in the absence of all other causes. Whereas causal support evaluates whether or not a causal relationship actually exists and calculates the strength of that relationship. To do this, causal support estimates the evidence for a graphical model with a link between *C* and *E* against one without a link. For example, let us consider the graphs denoted by Graph 0 and Graph 1 in Fig S1D (adapted from [33]). There are three variables in each graph: cause *C*, effect *E* and background cause B. In Graph 0, *B* causes *E*, but *C* has no relationship to either *B* or *E*. In Graph 1, both *B* and *C* cause *E*. While calculating ΔP and causal power Graph 1 is assumed, whereas causal support compares the structure of Graph 1 to that of Graph 0. Causal support is defined as the evidence provided from data D in favor of Graph 1, P(D | Graph 1), over Graph 0, P(D | Graph 0), which can be calculated by the following equation:

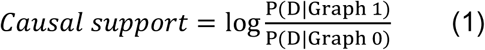

We calculated causal support using freely available Matlab code from [33]. ΔP and causal power were calculated using the following formulas:

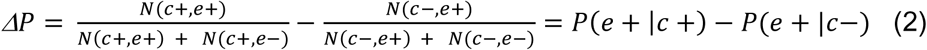

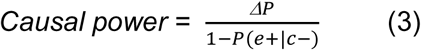

We compared these three measures of causal relationship with the transitional probability (i.e., a comparison between causation and correlation). We calculated the transitional probability of each greeble (V) given prior presentation of its paired sound (A), using the equation below:

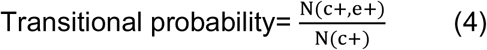

For each subject, we used the causal relationship value of each condition (HP, MP and NP) at the end of a testing phase as the starting values of the next testing phase. For example, the starting values of causal relationship for the “under drug” testing phase were equal to the causal relationship values at the end of the “pre/baseline” testing phase. Similarly, the starting values for the “after recovery” testing phase were equal to the causal relationship values at the end of the “under drug” phase (Fig S2A and B). We tested if subjects (i) regained access to already learned and stored predictive information, after they recovered from ketamine dosing, or (ii) re-learned the predictive information. We mimicked hypothesis (ii) by forcing the starting values of causal relationship (causal power here) “after recovery” to be zero (Fig S2C) instead of starting values equal to the causal relationship at the end of the “under drug” phase (hypothesis (i); Fig S2B).

#### Pharmacology Experiments

To manipulate participants’ predictions, we administered two drugs, ketamine and dexmedetomidine, each on a separate day, with at least one month intervening. The fixed order of dexmedetomidine in the first session, and ketamine in the second session, for the first 12 ketamine subjects was IRB-imposed in their consideration of safety profiles of the different medications (registered on NCT03284307); whereas the final three ketamine subjects were not administered dexmedetomidine in a prior session. All subjects were healthy and aged between 18-40 years old without contraindication to study drugs. We also acquired EEG data throughout pharmacology experiments (see section “EEG Recording” below), to measure electrophysiological activity during predictive coding. A typical pharmacology experiment consisted of three segments: (a) pre-drug baseline, (b) under drug influence (dexmedetomidine or ketamine), and (c) after recovery. During the pre-drug baseline, participants performed the learning phase (200 trials), then the first testing phase (400 trials). Under drug influence, participants performed the second testing phase (200 trials). Under ketamine, participants also performed a third testing phase (200 trials; see ketamine dosing for details). After recovery, they performed the last testing phase (400 trials). As a drug control, we tested three participants during saline administration. The saline results were similar to the pre-drug baseline. Due to a protocol-limited maximum time under drug influence, and the need to acquire sufficient M/NM trials for EEG analyses, the pharmacology experiments did not include inversion trials. All other aspects of the task in pharmacology experiments were the same as that in psychophysics experiments. For each of the two pharmacology experiments involving a particular participant, we used three new greeble name and image pairs to rule out any possible contribution of long-term memory.

#### Dexmedetomidine Dosing

We intravenously administered a 0.5 mcg/kg bolus over 10 minutes, followed by 0.5 mcg/kg/h infusion (MedFusion4000 pump at mcg/kg/hr). Participants performed the testing phase (under drug influence) during this infusion time, corresponding to stable drug levels according to the pharmacokinetic model for dexmedetomidine by Hannivoort et al. [83]. We targeted a plasma concentration of dexmedetomidine that is associated with mild sedation (modified observer’s assessment of alertness/sedation (OAA/S) of 4 [84]) to control for non-specific sedative effects, including hyperpolarization-activated cyclic nucleotide channel (HCN-1) effects [85]. The actual sedation achieved was on average slightly deeper than anticipated (modified OAA/S median 3, IQR 2) with a mean plasma concentration of 0.8 (SD 0.33) ng/ml [84].

#### Ketamine Dosing

In initial experiments, we tested two doses of ketamine to target the lowest plasma concentration that would modulate NMDARs in the relevant concentration range (<1microMolar [86]). The first dose corresponded to intravenously administered 0.25 mg/kg ketamine over 10 minutes, followed by 30 mg/h infusion, corresponding to 0.4 microMolar (Rugloop software using a Harvard 22 pump). Prior to testing subjects, lack of nystagmus and visual disturbance was confirmed in all participants. Participants performed the testing phase (under drug influence) during this infusion time, corresponding to stable drug levels according to the pharmacokinetic model for ketamine by Domino et al. [87, 88]. A second dose was tested with a second bolus of 0.25 mg/kg ketamine over 10 minutes, followed again by 30 mg/h infusion. Testing again was completed once stable plasma concentrations of approximately 0.8 microMolar were achieved. Ketamine blocked predictions at this second level of ketamine dosing, equating to a minimum plasma concentration of 0.2 μg/ml; range tested: 0.2-0.3 μg/ml. As we found the effective ketamine dose to be 0.2 μg/ml in our first seven subjects, we targeted that plasma concentration for our remaining subjects. We report data for the 0.2 μg/ml dosing in the manuscript. Three subjects were excluded from “after recovery” testing due to vomiting.

#### Monitoring

Subjects were monitored during drug exposure according to the American Society of Anesthesiologists guidelines, including electrocardiogram, blood pressure and oxygen saturation. We monitored arousal level using the modified OAA/S scale [89].

#### EEG Recording

We performed high-density EEG recordings using a 256 channel system (including NA 300 amplifier; Electrical Geodesics, Inc., Eugene, OR). After applying the EEG cap with conductive gel (ECI Electro-Gel), we adjusted electrodes so that the impedance of each electrode was within 0-50 kiloohms. We checked electrode impedance before the experiment started, and again before drug administration. Using Net Station, we sampled EEG signals at 250 Hz and, off-line, bandpass filtered between 0.1 Hz and 45 Hz.

#### EEG Preprocessing

We combined pre-drug baseline data from both ketamine and dexmedetomidine experiments (baseline RT data showed similar results for both drugs) but “under drug” analyses were performed separately for each drug. We performed offline preprocessing and analysis using EEGLAB [90]. First, we extracted data epochs -1,500 ms to 3,000 ms relative to the onset of the sound and -3000 ms to 800 ms relative to the onset of the image, for each trial. We then visually inspected each epoch and excluded noisy trials (around 5% of the total trials). Next, we performed Independent Component Analysis (ICA) using built-in functions of EEGLAB (pop_runica.m) and removed noisy components through visual inspection. We excluded three dexmedetomidine subjects from further analysis– due to very noisy EEG data, which after cleaning left too few trials for analysis (conditions with <10 trials). Finally, we performed channel interpolation (EEGLAB function, eeg_interp.m, spherical interpolation) and re-referenced to the average reference.

#### Time-Frequency Decomposition

To investigate changes in EEG spectral content, we performed time-frequency decompositions of the preprocessed data in sliding windows of 550 ms using Morlet wavelets, whose frequency ranged from 5 Hz to 45 Hz in 40 linearly spaced steps. Power for each time-frequency point is the absolute value of the resulting complex signal. We dB normalized power (dB power = 10*log10[power/baseline]) to the pre-stimulus baseline, starting 700 ms before sound onset (i.e., baseline calculated using sliding 550 ms-long wavelets, starting with the wavelet positioned from 700-150 ms before sound onset (centered on 425 ms before sound onset) and avoiding the sound-evoked response). For drift-diffusion model analysis, we calculated power spectral density and performed divisive baseline correction for each trial.

#### Electrode Selection

We used a data-driven approach, orthogonal to the effect of interest, to select the electrodes of interest based on the task EEG data. In the first step, we averaged power across all electrodes (aligned to image onset) and all sounds/greeble names. This revealed increased alpha power (8-14 Hz) during the delay period compared to baseline (Fig S3B). In the second step, we selected the electrode clusters that showed significant change in alpha power during the -925 ms to -275 ms time window pre-image onset, compared to baseline. This time window ensured that our delay period did not overlap with the sound or image. After cluster-based multiple comparisons correction (*6*), four different clusters showed significant modulation in alpha power during the delay period [and we selected one electrode (and its four nearest neighbors) in each cluster that showed the greatest modulation, for consistency across clusters; Fig 3A]: (i) right frontal (RF electrodes 4, 214, 215, 223, 224); (ii) left central (LC electrodes 65, 66, 71, 76, 77); (iii) right central (RC electrodes 163, 164, 173, 181, 182); and (iv) occipital (OC electrodes 117, 118, 119, 127, 129). Additionally, we found that only alpha power at the RF cluster significantly correlated with the predictive value of sounds.

#### Alpha Power Calculation

For each condition, HP, MP and NP, we averaged alpha power over all five electrodes in a cluster, to calculate the average alpha power of each cluster. To best capture the delay period activity just prior to image onset, we calculated mean alpha power between -625 to -275ms (as the wavelet window is centered around each time point, the power estimate before -625 ms and after -275 ms may contain auditory and visual stimulus-related responses, respectively) for each trial aligned to image onset. We also calculated mean delay period alpha power between 600 to 1,225 ms for each trial aligned to sound onset. To link EEG power spectral density to behavior using our drift-diffusion model analysis (see section HDDM), we calculated single-trial baseline-corrected (divisive normalization) alpha band power, aligned to image onset.

#### Source Space Analysis

We used FieldTrip’s beamforming technique to localize sources of the sensor level alpha activity [36]. This technique uses an adaptive spatial filter to estimate activity at a given location in the brain. We used a source model defined in MNI space for all subjects. Across all sound cues, we used the Dynamic Imaging of Coherent Sources (DICS) [91] algorithm to beamform the delay period alpha activity (window 0-900 ms before image onset). We then calculated the average source power for each region of interest (ROI) of the AAL atlas [92]. We selected ROIs that showed significant change in source power during the delay period (prior to image onset) compared to baseline (P < 0.025, Bonferroni corrected across all cortical AAL ROIs).

#### Granger Causality

We performed non-parametric spectral Granger causality analyses at the source level. Because there were no significant frontal ROIs in the left hemisphere, we restricted our analyses to the right hemisphere only. We used a covariance window of 0-900 ms prior to image onset and the Linearly Constrained Minimum Variance (LCMV) algorithm [93] to generate virtual time series for significant frontal and temporal ROIs. We averaged across the significant frontal ROIs (Superior Frontal Gyrus, medial; Superior Frontal Gyrus, dorsolateral; Middle Frontal Gyrus); and there was only one significant temporal lobe ROI (Inferior Temporal Gyrus). We calculated non-parametric spectral granger causality between the (average) frontal and temporal ROIs using the ft_connectivityanalysis function in FieldTrip for the stable window starting 900 ms prior to image onset [94, 95]. We found granger causal influence from right frontal to temporal cortex correlated with the predictive value of sounds only in the beta frequency (15-30 Hz) band.

#### Hierarchical Drift-Diffusion Model (HDDM)

We used a drift-diffusion model (DDM) [96], where there are two possible choices (correct/incorrect responses of the match trials) in our predictive coding task. According to this model, decision-making involves the accumulation of evidence (drift process) from a starting point to one of two (upper or lower) thresholds, representing the choices. The accumulation rate is known as the drift rate, *v*; and the starting point can be biased towards one of the choices (in our study, by the predictive value of the sound), reflected in a bias parameter, z. We used HDDM software (http://ski.clps.brown.edu/hddm_docs/) [29] for hierarchical Bayesian estimation of the parameters of the drift-diffusion model. Particularly with fewer trials per condition, this method has been shown to provide more reliable estimates of parameters and is less susceptible to outliers [96] than more traditional approaches to DDMs [97, 98].

To directly link the causal relationship between the sound and its paired greeble image to behavior and drift-diffusion parameters, we included the estimates of causal relationship and transitional probability as predictor variables of the bias, z, of the model. That is, we estimated posterior probability densities not only for basic model parameters, but also the degree to which these parameters are altered by variations in the psychophysical measures (ΔP, causal power, causal support and transitional probability). In these regressions, the bias parameter is given by, *z*(*t*) = *β*_0_ + *β*_1_*CP*(*t*), where CP is either ΔP, causal power, causal support or transitional probability, *β*_0_ is the intercept, and t is the trial number. Here, the slope, *β*_1_, is weighted by the value of the psychophysical measure on that trial. The regression across trials allows us to infer how the bias changes depending on the psychophysical measure. For example, if these psychophysical measures are positively correlated to bias, then increased causal strength or transitional probability will yield faster RTs. We fit four different versions of the model: (i) ΔP Model, where bias was estimated from the ΔP and updated after each trial; (ii) Causal Power Model, where bias was estimated from the causal power and updated after each trial; (iii) Causal Support Model, where bias was estimated from the causal support and updated after each trial; and (iv) Transitional Probability Model, where bias was estimated from the transitional probability and updated after each trial. We also modeled our data where drift rate varied according to causal strength or transitional probability. We used the Deviance Information Criterion (DIC) for model comparison [99]. The DIC is a measure of model fit (i.e., lack thereof) with a penalty for complexity (i.e., the number of parameters used to fit the model to the data) [100]. Models with lower DIC are better models. Models where bias was manipulated instead of drift rate had significantly lower DIC. For the rest of our analyses, we used models where bias varied with causal strength or transitional probability and drift rate was kept constant at the default values, as this yielded the lowest DIC.

We used Markov Chain Monte Carlo chains with 20,000 samples and 5000 burn-in samples for estimating the posterior distributions of the model parameters. We assessed chain convergence by visually inspecting the autocorrelation distribution, as well as by using the Gelman-Rubin statistic, which compares between-chain and within-chain variance. This statistic was near 1.0 for the parameters, indicating that our sampling was sufficient for proper convergence. We analyzed parameters of the best model (model with lowest DIC) using Bayesian hypothesis testing, where the percentage of samples drawn from the posterior fall within a certain region (e.g., > 0). Posterior probabilities ≥ 95% were considered significant. Please note that this value is not equivalent to p-values estimated by frequentist methods, but they can be coarsely interpreted in a similar manner.

All the model comparisons were estimated on the psychophysics data as these had the greatest number of trials per condition. This ensures robust estimation of the best model. The best-fitting model was then used to analyze data from different conditions: pre-drug baseline, under dexmedetomidine, under ketamine and recovery from ketamine. To directly link EEG power spectral density to behavior and drift-diffusion parameters, we used the HDDM, but now included the right frontal cluster power estimate (aligned to image onset) in the alpha band as the predictor variable of the bias, *z*, of the model; i.e., in the regression equation above, CP was now alpha power.

#### Event-related Potentials

We used ERPLAB (https://erpinfo.org/erplab) [101] to run event-related potential (ERP) analysis. First, we cleaned epoched data aligned to the sound onset for auditory ERPs using the pop_artmwppth.m function of ERPLAB with a moving window of 200 ms (2-3% of trials for each subject were excluded). We then averaged over trials to generate an average ERP for each subject. We used 200 to 0 ms prior to stimulus onset as baseline. Based on previous literature[102, 103], we chose channel 9 (Cz electrode) for exemplar auditory ERPs in figures.

#### EEG Signal Decoding

We used two different machine learning approaches in this study: (i) a Support Vector Machine (SVM) model to classify HP, MP and NP trials after sound onset, with EEG power spectral density as input; and (ii) a recurrent neural network (RNN) to classify V1, V2 and V3 trials relative to image onset, with “raw” EEG time series data as input.

We used the power spectral density of EEG signals across four frequency bands, theta (5-7 Hz), alpha (8-14Hz), beta (15-30 Hz) and gamma (31-40 Hz), for the first decoding analysis. We calculated the sum of squared absolute power in each frequency band for each electrode cluster (RF, LC, RC and OC), thus generating 16 features for each trial as the input dataset. Using Scikit-learn [104] implemented in Python, for each 20 ms time bin, we trained a SVM model to classify the EEG data into three classes: highly predictive (HP), moderately predictive (MP) and not-match predictive (NP). We denote *y_i_*(*t*) ∈ {0,1,2} as the identifier of the condition at time bin *t*, where 0, 1, and 2 denotes HP, MP and NP respectively. The SVM classifier is implemented by a nonlinear projection of the training data **x**(*t*) feature space *X* into a high dimensional feature space *F* using a kernel function *ϕ*. So with *ϕ* : 𝓧→𝓕 being the mapping kernel, the weight vector **w** can be expressed as a linear combination of the training trials and the kernel trick can be used to express the discriminant function as

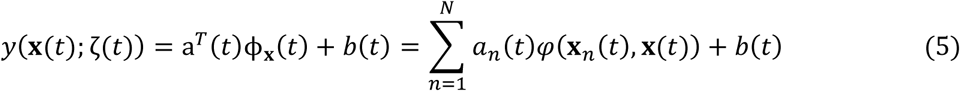

where *ζ*(*t*) = {*a*(*t*), *b*(*t*)} is the new parameter at time bin *t* with *a*(*t*) and b(*t*) as weights and biases of the mapped features space 𝓕. We used the radial basis function (RBF) kernel that allows nonlinear decision boundary implementation in the input space. The RBF kernel holds the elements

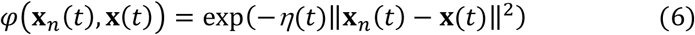

Where *η*(*t*) is a tunable parameter at each time bin. Model hypermeters consisting of regularization penalty (*C*(*t*)) and *η*(*t*) were selected by grid search through 10-fold cross validation. F-score at each time bin and for each label was calculated as

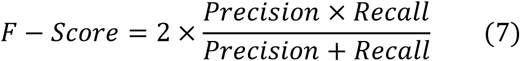

where

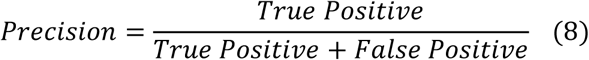

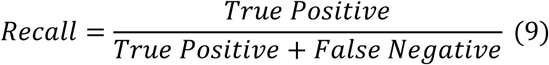

As we penalized the mapped weights of the classifier at each time bin, we used normalized absolute values of the weights as a measure to deduce each feature’s contribution to classify the outputs.

Second, we used a RNN with many-to-many architecture to decode visual stimuli using the EEG time domain signals from the four clusters of electrodes (RF, RC, LC, OC) mentioned above (20 features in total) in successive 20 ms time bins. The proposed RNN consists of a Bidirectional long-short term memory (BiLSTM) layer followed by an attention layer, a fully connected with dropout layer and a softmax layer as output. Considering x(t) as the input at time t, the output of the BiLSTM forward path is calculated as follow:

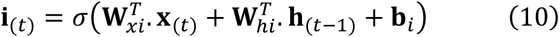

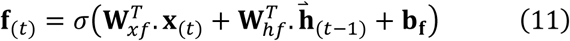

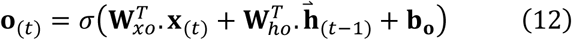

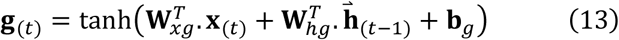

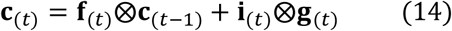

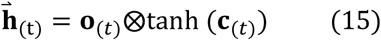

Where **i**_(*t*)_, **f**_(*t*)_, **c**_(*t*)_, **o**_(*t*)_ are the input gate, forget gate, cell gate and output gate respectively, 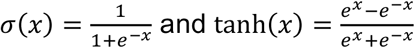, and the output of the backward path is 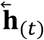. The attention layer output can be calculated as follows:

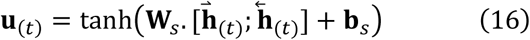

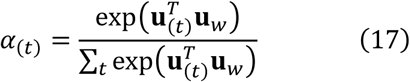

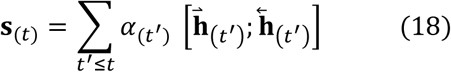

the output of the attention layer was fed to a fully connected layer with dropout followed by a softmax layer which results in class conditional probability as equation (19) specifies.

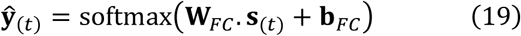

All BiLSTM, attention and FC layers weights and biases will be updated through backpropagation in time. Hyperparameters of the model including the number of hidden units for BiLSTM, Attention and FC layers, learning rate, dropout probability, learning rate of stochastic gradient descent, etc., are optimized via a grid search. The decoding accuracy is calculated with 10-fold cross validation with 60% of trials as training set and 40% as test set by shuffling and stratifying to avoid accuracy bias due to imbalance classes.

To find out how well the model can be generalized in time, we tested the trained model until time *t* with the data at all time points using the stored parameters of the above RNN model. The results of this analysis is used to see whether information about the visual stimulus can be decoded prior to stimulus presentation.

To gain insight on how well the model decodes each visual stimulus during the delay period, we trained the model on data at 100 ms post-visual stimulus onset and calculated the F-score for each class (V1, V2, V3) under all conditions (before drug, under ketamine and DEX).

Using the same model (trained on data 100ms after visual stimulus onset), we looked at the proportion of each auditory stimulus (HP, MP and NP) associated with the correctly decoded trials at 100 ms before visual stimulus onset.

### STATISTICAL ANALYSIS

We performed statistical analysis of trials from testing phases and used the learning phase only to confirm that the participants learned the correct associations. To only include the trials where the causal power of HP, MP and NP trials have differentiated (Fig 1G), we excluded the first 50 trials of the testing phase in psychophysics experiments and the pre-drug baseline. We used all available trials for the testing phase of “under drug” and “after recovery”, as the causal power of HP, MP and NP trials were already differentiated from the beginning (Fig S2A and B). For RT analysis, we excluded RTs more/less than the mean +/- 3SD for each subject, and we used correct match trials. For delay period EEG analyses, we used both correct match and non-match trials.

Because we had clear, a priori predictions about the effects of the different conditions in this study, i.e., the effect of prediction and the effects of the drug conditions, we could apply contrast analysis using linear mixed effects models (LMEs). Our study conformed to the guidelines set out by Ableson and Prentice [30]; i.e., we included the contrast of interest along with a paired, orthogonal contrast and infer significance only when our statistical tests showed that effects were significant for the contrasts of interest and not for the orthogonal contrasts.

EEG data were analyzed using contrast analysis with a LME where the predictive value of sound (HP, MP, NP; varying within subject) and drug condition (before ketamine, under ketamine, under dexmedetomidine; varying within subject) served as independent variables. For prediction, we used a linear (−1, 0, 1) contrast as our contrast of interest, and a quadratic (−1, 2, -1) contrast as the orthogonal contrast in the analysis. For drug condition, we used a quadratic (−1, 2, -1) contrast as our contrast of interest, and a linear (−1, 0, 1) contrast as the orthogonal contrast in the analysis. Mathematically, our LME can be written as equation 20 below.

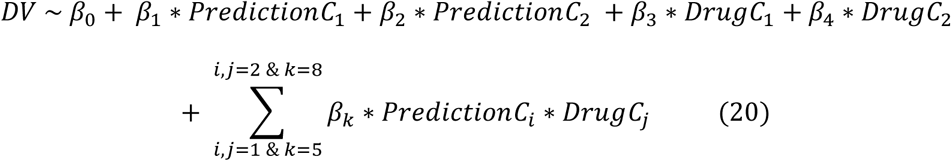

Here, DV is the dependent variable (RF alpha power or average GC at beta frequency), *PredictionC*_1_ is the contrast of interest modelling sounds’ predictive value, *PredictionC*_2_ is the orthogonal contrast for sounds’ predictive value, *DrugC*_1_ is the contrast of interest modelling drug condition, *DrugC*_2_ is the orthogonal contrast for drug condition. Terms after the summation notation in equation 20 correspond to interactions between sounds’ predictive value and drug condition (all possible interactions, i.e. permutations of contrasts, included).

We used a similar LME with prediction (HP, MP, NP; varying within subject) and drug condition (under ketamine, before drug, under DEX, after recovery; varying within subject) as independent variables to analyze RT data. We used a linear (HP, MP, NP: -1, 0, 1) contrast as our contrast of interest to model RT, and a quadratic (HP, MP, NP: -1, 2, -1) contrast as the orthogonal contrast in the analysis. For drug condition, we used a four-level (under ketamine, before drug, under DEX, after recovery: 3, -1, -1, -1) contrast as our contrast of interest, and two orthogonal contrasts (under ketamine, before drug, under DEX, after recovery: 0, 0, -1, -1 and 0, 2, -1, -1) in the analysis. Both main effects of sounds’ predictive value and drug as well as their interaction were included. We used contrast analysis to model our hypotheses. Mathematically, our LME can be written as equation 21 below.

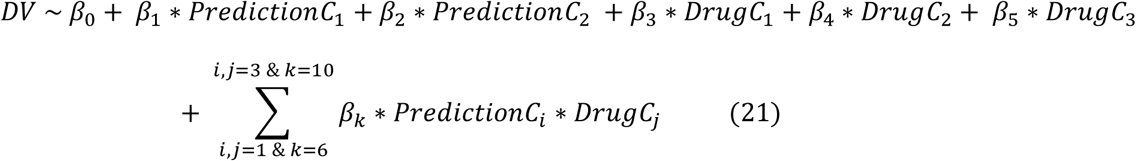

Here, DV is the dependent variable (RT or accuracy), *PredictionC*_1_ is the contrast of interest modelling sounds’ predictive value, *PredictionC*_2_ is the orthogonal contrast for sounds’ predictive value, *DrugC*_1_ is the contrast of interest modelling drug condition, *DrugC*_2_ and *DrugC*_3_ are the orthogonal contrasts for drug condition. Terms after the summation notation of equation 21 correspond to interactions between sounds’ predictive value and drug condition (all permutations of two-way interactions included).

To investigate auditory ERPs, we ran non-parametric permutation tests using the *ft_timelockstatistics.m* function from Fieldtrip [36]. For before and during ketamine conditions, we looked for differences in ERP between sound cues across 50 ms to 300 ms latency from sound onset. We randomly shuffled condition labels 10,000 times. Alpha value was set to 0.05 for all comparisons.

We used repeated measures ANOVA (Holm-Sidak corrected p-values) to test the significance of F-scores for the three differentially predictive conditions (HP, MP, NP) as well as the significance of each feature’s contribution in output classification.

To test the significance of the cross-temporal decoding results, we performed a paired t-test between the 20-fold cross validated and 100 times resampled accuracy at each pixel and randomly permuted output labels model at the same pixel. The resulting p-values then were corrected for multiple comparisons (Holm-Sidak corrections).

## Acknowledgements

We thank G. Lupyan, R.A. Pearce and B.R. Postle for useful discussions. NIH grants R01MH110311 (Y.S.), R01NS117901 (Y.S and R.S.), K23AG055700 (R.S.) and R01AG063849 (R.S.) supported this work

---

**Fig. S1.**
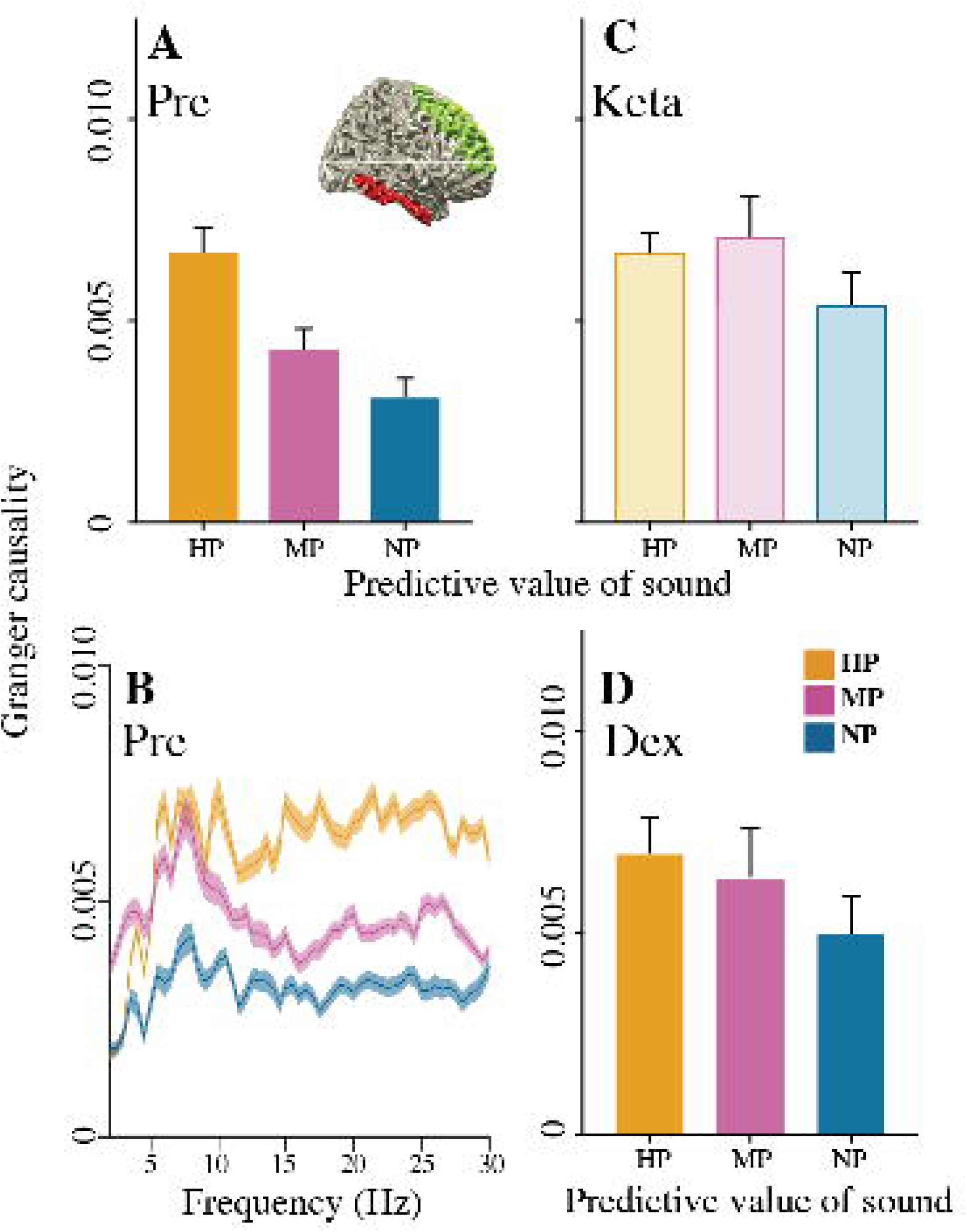
Fast RTs to predictive sounds not due to speed-accuracy trade off. Related to Fig 1. Population average (25 subjects from psychophysics experiment) (A) RT and (B) accuracy for highly predictive (HP), moderately predictive (MP), and not-match predictive (NP) sounds. (C) Posterior probability density of β_1_ of hierarchical drift diffusion model from psychophysics experiment. (D) Directed graphs for calculating causal support. In graph 0, B causes E, but C has no relationship to either B or E. In graph 1, both B and C cause E (adapted from [33]). (E) Transitional probability (TP) of an exemplar subject across time for pre-ketamine (Pre) testing. Each data point corresponds to the TP at a particular trial. (F) Causal power (CP) of an exemplar subject across time for pre-ketamine (Pre) testing. Each data point corresponds to the CP at a particular trial.

**Fig S2.**
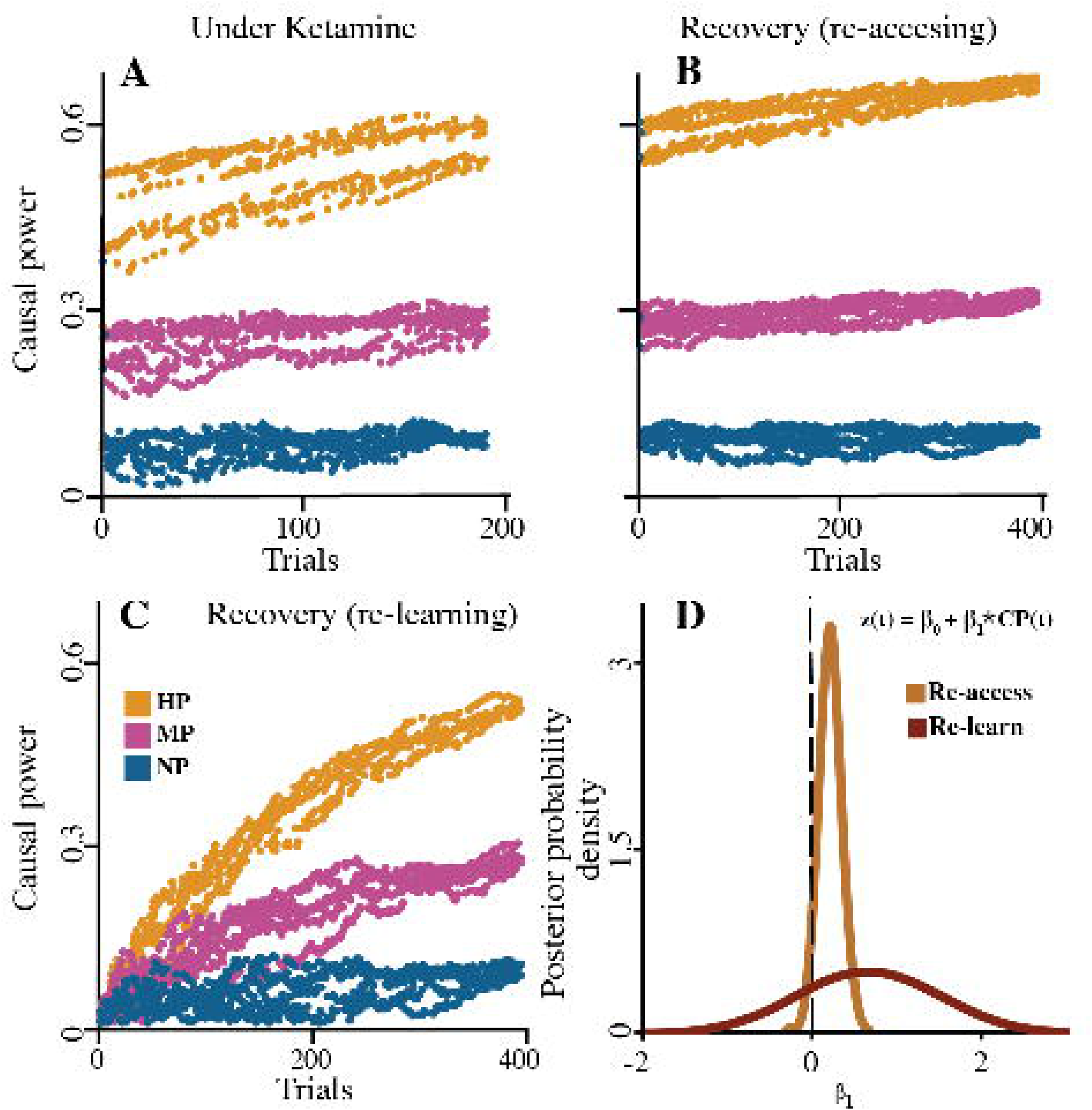
Ketamine blocked access to predictive information. Related to Fig 1. (A) Population causal power values for subjects under ketamine for highly predictive (HP), moderately predictive (MP), and not-match predictive (NP) sounds. (B) Population causal power values after recovery when subjects regained access to predictive information. (C) Population causal power values for hypothetical condition where subjects re-learned predictive information. (D) Positive β_1_ of hierarchical drift diffusion model for first 30 trials after recovery from ketamine, when subjects re-accessed predictive information (P{β_1α_>0}=0.03). In contrast, the β_1_ for re-learning was not greater than zero (P{β_1α_>0}=0.20).

**Fig S3.**
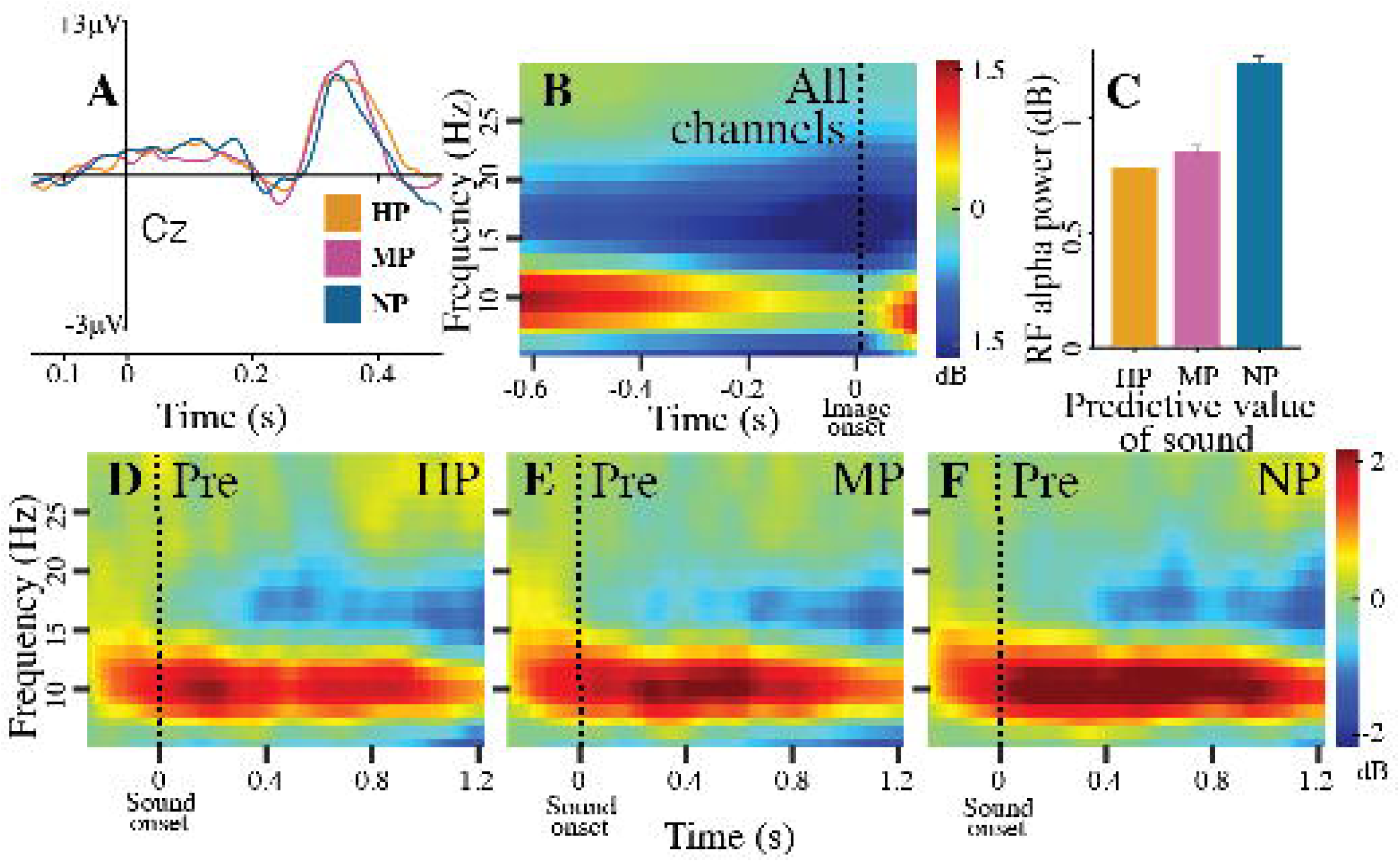
Early sensory processing differences cannot account for prediction strength as all three sounds generated similar auditory ERPs. Related to Fig 2. (A) Exemplar auditory ERPs at Cz electrode for highly predictive (HP), moderately predictive (MP) and not-match predictive (NP) sounds. (B) Time-frequency plot of power averaged over all electrodes and all trials. Power calculated in 0.55 s sliding windows, with window at 0 s representing interval - 0.275 s to +0.275 s. Plot aligned to image onset. (C) Population average RF alpha power (+SE) during delay period; aligned to sound onset. Time-frequency decomposition of right frontal electrode cluster (RF) before drug administration (Pre) for HP (D), MP (E) and NP (F) sounds. Power calculated in 0.55 s sliding windows, with window at 0 s representing interval -0.275 s to +0.275 s. Plots aligned to sound onset.

**Table S1.** 2 X 2 contingency table for each sound. Related to Fig 1. Generic contingency table for all sounds in black. N(c^+^,e^+^) represents the number of trials in which the effect occurs in the presence of the cause, N(c^-^,e^+^) represents the number of trials in which the effect occurs in the absence of the cause, N(c^+^,e^-^) represents the number of trials in which the cause occurs but not the effect, and N(c^-^,e^-^) represents the number of trials in which the cause and effect are absent. ‘Cause’ used in statistical sense. In green, example contingency table for sound A1 where N(c+,e+) are the number of trials V1 follows A1 (A1-V1); whereas N(c-,e+) would be the number of trials V1 follows A2 or A3 (A2-V1 or A3-V1) and so on.

| | Effect Present ($e^+$)<br>*Effect Present (V)* | Effect Absent ($e^-$)<br>*Effect Absent* |
| --- | --- | --- |
| Cause Present ($c^+$)<br>*Cause Present (A)* | $N(c^+, e^+)$<br>*A1-V1* | $N(c^+, e^-)$<br>*A1-V2, A1-V3* |
| Cause Absent ($c^-$)<br>*Cause Absent* | $N(c^-, e^+)$<br>*A2-V1, A3-V1* | $N(c^-, e^-)$<br>*A2-V2, A2-V3, A3-V2, A3-V3* |

